# Abscisic Acid Increases Hydrogen Peroxide in Mitochondria to Facilitate Stomatal Closure

**DOI:** 10.1101/2022.01.11.475946

**Authors:** Anthony E. Postiglione, Gloria K. Muday

## Abstract

Abscisic acid (ABA) drives stomatal closure to minimize water loss due to transpiration in response to drought. We examined the subcellular location of ABA increased accumulation of reactive oxygen species (ROS) in guard cells that drive stomatal closure. ABA-dependent increases in fluorescence of the generic ROS sensor, dichlorofluorescein (DCF), were observed in mitochondria, chloroplasts, cytosol, and nuclei. The ABA response in all these locations were lost in an ABA-insensitive quintuple receptor mutant. The ABA-increased fluorescence in mitochondria of both DCF and an H_2_O_2_-selective probe, Peroxy Orange 1 (PO1), colocalized with Mitotracker Red. ABA treatment of guard cells transformed with the genetically-encoded H_2_O_2_ reporter targeted to the cytoplasm (roGFP2-Orp1), or mitochondria (mt-roGFP2-Orp1), revealed H_2_O_2_ increases. Consistent with mitochondrial ROS changes functioning in stomatal closure, we found that guard cells of a mutant with mitochondrial defects, *abo6*, have elevated ABA-induced ROS in mitochondria and enhanced stomatal closure. These effects were phenocopied with rotenone, which increased mitochondrial ROS. In contrast, the mitochondrially targeted antioxidant, MitoQ, dampened ABA effects on mitochondrial ROS accumulation and stomatal closure in Col-0 and reversed the guard cell closure phenotype of the *abo6* mutant. ABA-induced ROS accumulation in guard cell mitochondria was lost in mutants in genes encoding Respiratory Burst Oxidase Homolog (RBOH) enzymes and reduced by treatment with the RBOH inhibitor VAS2870, consistent with RBOH machinery acting in ABA-increased ROS in guard cell mitochondria. These results demonstrate that ABA elevates H_2_O_2_ accumulation in guard cell mitochondria to promote stomatal closure.

**One sentence summary:** Genetically encoded biosensors and chemical probes revealed ABA-dependent increases in hydrogen peroxide, a reactive oxygen species with signaling activity, in guard cell cytoplasm and mitochondria.

## Introduction

Drought stress negatively impacts plant growth due to excess water loss, which is a growing concern for crop yields as a result of the changing global climate (Fahad et al., 2017). Stomatal closure reduces excess water loss but also limits CO_2_ entry, thereby negatively impacting the photosynthetic rate (Lamaoui et al., 2018). Due to this tradeoff, stomatal aperture must be tightly controlled (Nilson and Assmann, 2007). Reduction in guard cell turgor to close stomata is mediated by the hormone abscisic acid (ABA), which signals in guard cells during states of decreased water availability (Xu et al., 2016; Li et al., 2017; Qi et al., 2018; Qu et al., 2018; Tõldsepp et al., 2018).

The binding of ABA to the PYR/PYL/RCAR family of soluble receptors initiates an ABA signaling cascade (Park et al., 2009). The ABA bound receptors form a complex with Clade A protein phosphatases type 2C (PP2Cs), which negatively regulate ABA signaling in the absence of the hormone (Hsu et al., 2021). Formation of this complex inhibits PP2C activity, releasing the negative regulation of the pathway (Ma et al., 2009; Park et al., 2009; Nishimura et al., 2010). Reduced phosphatase activity allows for increased phosphorylation of a variety of proteins including Sucrose nonfermenting Related Kinase 2 family members (SnRK2s) (Takahashi et al., 2020). Active SnRK2s can then further promote the signaling cascade through phosphorylation of a number of downstream targets including Respiratory Burst Oxidase Homologs (RBOH) enzymes, also called NADPH oxidase (NOX) enzymes (Sirichandra et al., 2009). Consistent with RBOH activation, ROS accumulation in guard cells following ABA treatment has been observed in multiple plant species (Pei et al., 2000; Kwak et al., 2003; Watkins et al., 2014; Watkins et al., 2017). These elevated ROS act as second messengers that lead to decreases in H^+^-ATPase activity and K^+^ uptake, while increasing efflux of K^+^, Cl^-^, and malate. This results in the reduction of guard cell turgor, and closure of the stomatal pore (Geiger et al., 2009; Jezek and Blatt, 2017; Demidchik, 2018; Klejchova et al., 2021).

ROS bursts resulting from RBOH activation have been characterized in plants in response to a myriad of developmental and environmental signals (Chapman et al., 2019; Martin et al., 2022). These enzymes function in the production of extracellular superoxide through the transfer of electrons from NADPH or FADH_2_ to molecular oxygen (Suzuki et al., 2011). Superoxide may be rapidly converted to H_2_O_2_ spontaneously or by enzymatic means via superoxide dismutases (Fukai and Ushio-Fukai, 2011). Extracellular H_2_O_2_ may then enter plant cells through plasma membrane localized aquaporins (Bienert et al., 2007; Tian et al., 2016; Rodrigues et al., 2017). The Arabidopsis genome encodes 10 RBOH family members (RBOHA-RBOHJ) with distinct expression patterns and functions that regulate a variety of developmental and cellular processes (Chapman et al., 2019). Genetic approaches have identified a role for RBOHF during ABA-induced stomatal closure (Kwak et al., 2003). Both the *rbohf* single mutant and the *rbohd/f* double mutant displayed a reduction in both ABA-driven ROS increases and stomatal closure as compared to wild-type guard cells (Kwak et al., 2003).

Insight into the subcellular localization of where ROS accumulates in guard cells after ABA treatment, as well as the type of ROS that are increased, are needed to fully understand how these molecules function in ABA signaling. Prior studies examining ABA-dependent increases in ROS accumulation have largely examined changes in fluorescence of dichlorofluorescein (DCF), the cleavage product of CM 2,7-dihydrodichlorofluorescein diacetate (CM H_2_DCF-DA), a cell permeable generic ROS sensor (Pei et al., 2000; Zhang et al., 2001; Kwak et al., 2003; An et al., 2016; Wu et al., 2017), which does not reveal which types of ROS are increased by ABA (Kalyanaraman et al., 2012; Winterbourn, 2014). Prior work has shown that the transcriptional responses downstream from ROS signals appear to hold a level of specificity not only based on what type of ROS is being sensed, but also where a given ROS is being generated (Gadjev et al., 2006). While redox changes in plants have been characterized in response to a myriad of environmental responses in multiple tissues, studies providing details on the type of ROS produced and where the ROS are accumulating are sparse. Therefore, there are still many questions remaining that require ROS detection methodology that can allow visualization of specific types of ROS in a particular subcellular compartment. While this is a difficult task given the highly reactive nature of these molecules, excellent advancements have been made in both microscopic resolution and ROS detection using chemical or genetically encoded sensors, with the roGFP2-Orp1 bioreporter providing significant insight into the localization of hydrogen peroxide (Dickinson et al., 2010; Winterbourn, 2014; Nietzel et al., 2019; Ugalde et al., 2021). Together these tools have provided the ability to gain better insight into this spatial accumulation of different ROS species in response to environmental stresses, hormone signaling and development.

This study examined how ABA affects the accumulation and localization of H_2_O_2_ within Arabidopsis guard cells during the ABA response and how each of these intracellular compartments contribute to total ROS changes that drive stomatal closure. The subcellular distribution of the general ROS sensor, DCF, was examined across multiple subcellular locations including the guard cell chloroplasts, cytosol, nuclei, and cytosolic puncta that we identified as mitochondria. To verify that the compartmentalized ROS changes were directly tied to ABA signaling, we examined these ROS changes in an ABA quintuple mutant. To determine if H_2_O_2_ increases in response to ABA, both Peroxy Orange 1, a chemical probe selective for H_2_O_2_, and the roGFP2-Orp1 genetically encoded hydrogen peroxide sensor, which was targeted to either the cytosol, mitochondria, or chloroplast, were used to provide insight into changes in this signaling ROS. Genetic and pharmacological approaches were also used to manipulate mitochondrial ROS production to test the role of this localized ROS in ABA-dependent stomatal closure. This combination of chemical, genetic, and imaging approaches reveals that ABA increases H_2_O_2_ in guard cell mitochondria and that ROS increases within this organelle play a necessary role in ABA-induced stomatal closure.

## Results

### ABA Signaling Drives Compartmentalized ROS Increases within Guard Cells

ABA-induced ROS increases within Arabidopsis guard cells, were verified through quantification of fluorescence intensity changes using a generic ROS-responsive fluorescent probe. We utilized CM 2,7-dihydrodichlorofluorescein diacetate (CM H_2_DCF-DA), which is a frequently utilized fluorescent chemical probe to monitor changes in ROS accumulation (Halliwell and Whiteman, 2004). CM H_2_DCF-DA diffuses across the plasma membrane, where it is trapped within the cell after cleavage by cellular esterases (Swanson et al., 2011). The probe is then converted to the highly fluorescent DCF upon oxidation by ROS. Whole leaves of Col-0 were excised in the morning, epidermal peels were prepared, and then covered with a stomatal opening solution for 3 hrs under white light to fully open stomata. This was followed by incubation with 20 μM ABA or a control treatment for 45 min. Treatments were then removed and leaf peels were incubated with CM H_2_DCF-DA for 15 min. Laser scanning confocal microscope (LSCM) images of guard cells with control or ABA treatment with the images of DCF fluorescence are shown directly and after conversion to lookup tables (LUT), which clarifies the range of fluorescence across these cells, with representative images shown in Supplemental Figure S1A. Whole stomata DCF fluorescence was recorded in 30 or more stomata per treatment and each individual value was normalized relative to the average signal intensity of control buffer treated stomata. This quantification confirmed that ABA significantly increased DCF fluorescence in guard cells during ABA-induced stomatal closure (Supplemental Figure S1B).

The images in Supplemental Figure S1 suggest that ABA increases DCF signal in a number of distinct subcellular locations. We therefore performed high resolution imaging of Col-0 guard cells in the presence and absence of 20 μM ABA for 45 min to identify the location within the cell that ABA signaling increased ROS. ABA treatment increased DCF fluorescence within multiple subcellular regions (Figure 1). The DCF signal increases in the chloroplast and nucleus were verified through spectral unmixing of the DCF signal from chlorophyll autofluorescence and DAPI fluorescence, respectively (Supplemental Figure S2A-B). DCF fluorescence is largely excluded from the vacuole, but there are increases in DCF signal after ABA treatment in the chloroplasts, cytosol, nucleus, and small cytosolic punctate structures (Figure 1A), which we identified as mitochondria in experiments described below. To determine how each of these subcellular regions contribute to overall ROS changes, we quantified increases in each of these locations. Following 45 minutes of 20 μM ABA treatment we observed the largest increases within the mitochondria and chloroplasts at 1.5-fold and 1.8-fold, respectively, as compared to control treated guard cells (Figure 1B-C). We also observed significant increases in other locations with a 1.3-fold ABA-induced increase within guard cell nuclei (Figure 1D), and a 1.4-fold increase in the cytosolic signal (Figure 1E).

**Figure 1.**
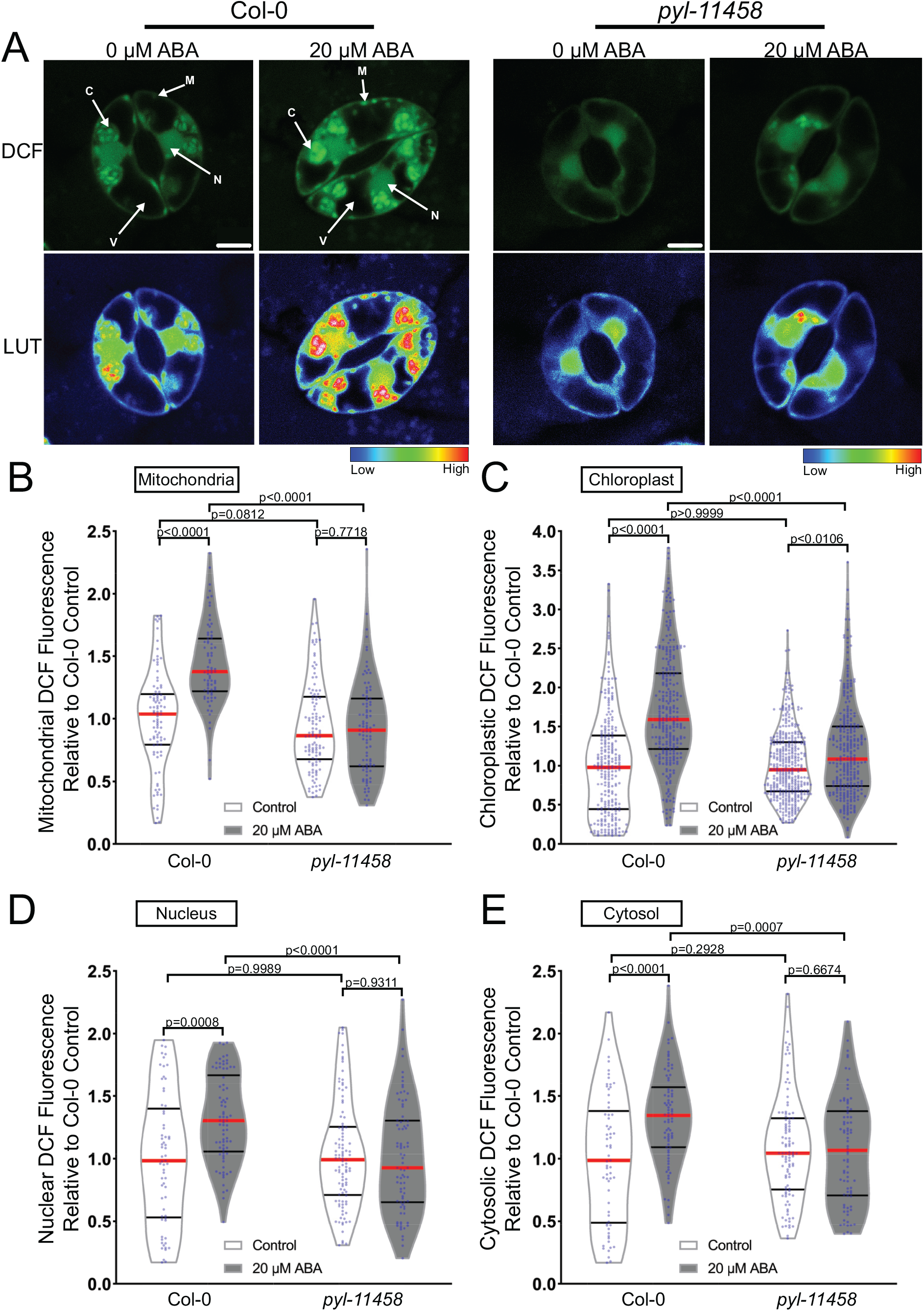
ABA treatment increases DCF fluorescence in multiple subcellular locations within Arabidopsis guard cells. A) Confocal micrographs of DCF fluorescence of guard cells in leaves from Col-0 or a quintuple ABA receptor mutant, *pyl-11458*, treated with control buffer or 20 μM ABA shown directly or after conversion to LUT. Subcellular compartments are indicated on each image (C: Chloroplast, N: Nucleus, M: Mitochondria, V: Vacuole). Scale bar: 5μm. Quantifications of DCF fluorescence in the B) mitochondria, C) chloroplasts, D) nucleus and E) cytosol with and without ABA treatment from three separate experiments (chloroplast n>100, nucleus n>48, cytosol n>48 and mitochondria n>56). All individual values were normalized to the average of the control treatment for Col-0 in each subcellular location and are displayed on the graph as blue dots with the median shown in red and lower and upper quartiles indicated in black. All p-values were calculated by two-way ANOVA followed by a Tukey’s post-hoc test from at least three separate experiments.

To determine whether these subcellularly localized ABA-induced ROS changes were downstream of the canonical ABA signaling pathway, we also examined the response in the ABA quintuple receptor mutant (*pyl-11458*). The DCF fluorescence in the mitochondria, nucleus, and cytoplasm were no longer significantly different between buffer and ABA treatment in the *pyl-11458* mutant. In the chloroplasts, the magnitude of the ABA-dependent increase was reduced relative to Col-0, although there was still a significant difference in response to ABA treatment (Figure 1B-E). These data suggest that ABA increased ROS in several intracellular locations within guard cells, however DCF cannot resolve which reactive oxygen species increases.

### ABA Increases H_2_O_2_ in Subcellular Regions Including Chloroplasts and Mitochondria to Induce Stomatal Closure

To ask whether hydrogen peroxide (H_2_O_2_) is the ROS that increases in response to ABA, we utilized the H_2_O_2_-selective chemical probe, Peroxy Orange 1, (PO1) (Dickinson et al., 2010) which is a membrane-permeable, boronate-based probe that becomes fluorescent upon irreversible oxidation by H_2_O_2_. The spectral profile of PO1 was isolated and unmixed from leaf auto-fluorescence, including chloroplasts, and the signal in the absence and presence of ABA is shown in Figure 2A. Treatment with 20 μM ABA significantly increased PO1 signal in the chloroplasts but did not result in a significant increase in the mitochondria (Figure 2B-C), nucleus, or cytosol (Supplemental Figure S3). To determine if higher concentrations of ABA were sufficient to stimulate H_2_O_2_ increases throughout guard cells, we increased concentration of ABA to 100 μM.

**Figure 2.**
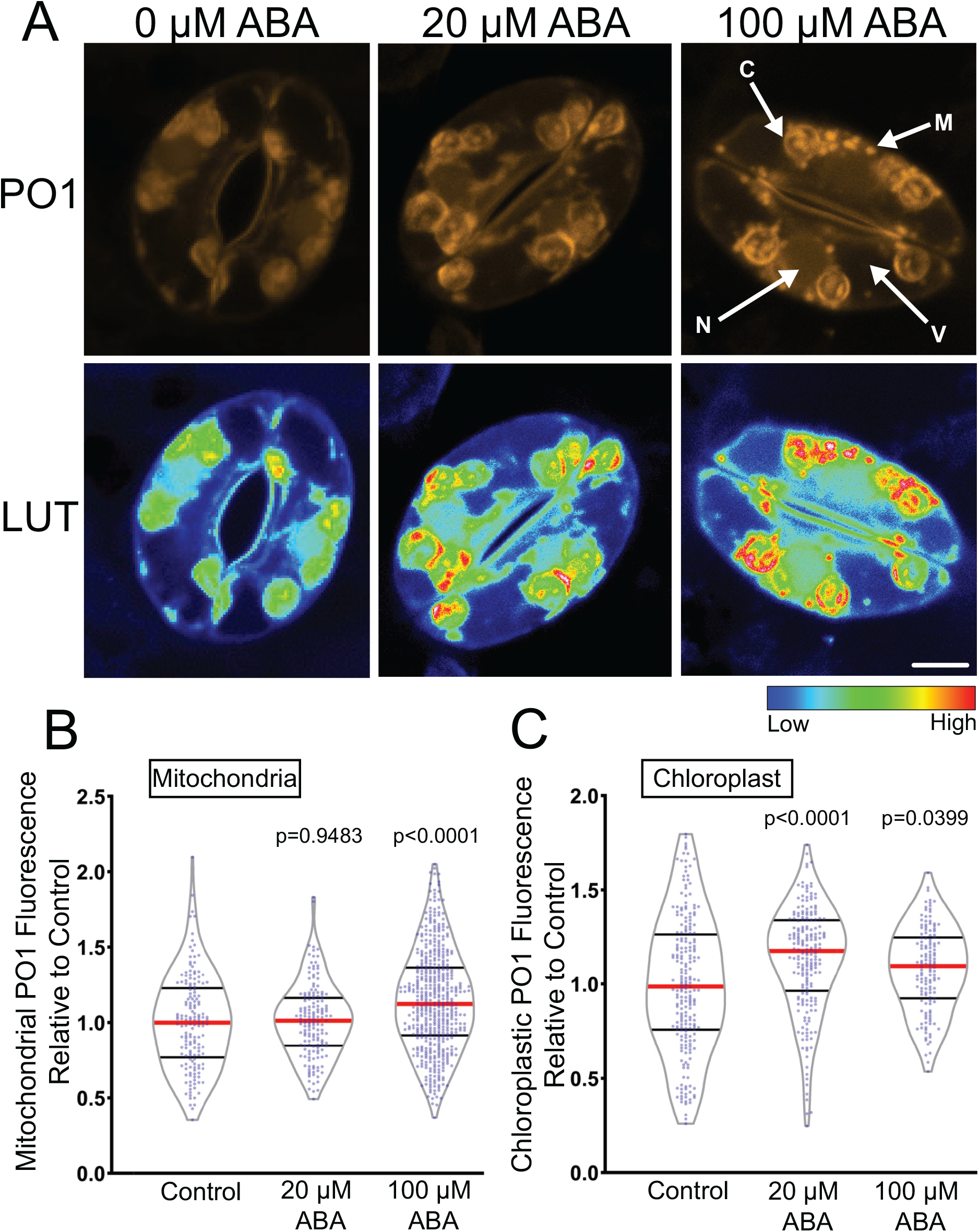
ABA increases fluorescence of Peroxy Orange 1 (PO1), a hydrogen peroxide selective dye. A) Confocal micrographs of PO1 fluorescence or PO1 signal converted to Lookup Tables (LUT) after treatment with 0, 20, or 100 μM ABA. Maximum intensity projections of full z-stacks are shown. Scale bar: 5μm. Quantifications of PO1 fluorescence in B) mitochondria and C) chloroplasts. All individual values were normalized to the average of the control treatment for each subcellular location and are displayed on the graph as blue dots with the median shown in red and lower and upper quartiles indicated in black. All p-values were calculated by one-way ANOVA followed by a Tukey’s post-hoc test from at least three separate experiments (mitochondria n>148 and chloroplast n>140).

Following treatment with 100 μM ABA, we observed a small but significant increase in PO1 signal within mitochondria (Figure 2B) as well as a significant increase in PO1 signal in chloroplasts (Figure 2C). However, PO1 signal in the cytosol and nucleus were once again not significantly different from control treatments (Supplemental Figure S3C-D). This suggests that PO1 may not be taken up by all organelles or that PO1 may not be sensitive enough to detect the H_2_O_2_ changes displayed in certain subcellular locations.

To verify that PO1 fluorescence is ROS responsive in the nucleus, we treated guard cells with 250 μM exogenous H_2_O_2_ (Supplemental Figure S4A-C). Treatment with exogenous H_2_O_2_ resulted in a 1.5-fold increase in nuclear PO1 fluorescence indicating that PO1 can be sufficiently taken up by Arabidopsis guard cell nuclei to detect large increases in H_2_O_2_ (Supplemental Figure S4C). The effect of this exogenous H_2_O_2_ treatment on stomatal closure was quantified in Supplemental Figure S4B. Treatment with exogenous H_2_O_2_ for 30 minutes was sufficient to close stomata to levels consistent with ABA-dependent closure. Together, these results suggest that H_2_O_2_ can function as the ROS to close stomata, but that ABA-induced increases in H_2_O_2_ in guard cells may need more sensitive tools than PO1 for their detection.

In the chloroplast, we observed ABA-induced PO1 accumulation into distinct structures within inner chloroplast compartments. However, two-dimensional images of maximum intensity projection make it difficult to discern if these dyes are associated with the chloroplast membrane, or internal to this organelle. Therefore, we created three-dimensional renderings of PO1 labeled guard cells to gain insight into chloroplast PO1 distribution (Supplemental Figure S5). These renderings show that bright PO1 regions span the chloroplast and are not just on the surface, revealing a complex ROS accumulation pattern within the chloroplast while the mitochondria are circular structures with uniform PO1 signal. The PO1 localized signal is similar in position and size to the accumulation of starch grains (Leshem and Levine, 2013), though we cannot currently rule out that the complex accumulation pattern of PO1 fluorescence in chloroplasts is due to increased dye sequestration in regions within this organelle.

### ABA Treatment Results in DCF Increases within Guard Cell Mitochondria

DCF and PO1 localized to small cytosolic punctate structures in addition to chloroplasts and nuclei. We used colocalization of chemical ROS probes with fluorescent organelle dyes and an organelle targeted GFP to ask whether these puncta are ROS producing peroxisomes or mitochondria (Figure 3). To evaluate whether cytosolic ROS puncta were peroxisomes, we examined an Arabidopsis transgenic line with a GFP tagged with a type 1 peroxisomal targeting signal (GFP-PTS1) (Ramón and Bartel, 2010) (Figure 3A). GFP-PTS1 and PO1 have emission peaks that can be spectrally unmixed. We used the Zen colocalization module to draw ROIs around cytosolic punctate structures that did not overlay on a chloroplast (Figure 3B). Although the GFP-PTS1 signal accumulated into punctate structures within the guard cell cytosol, they did not colocalize with the puncta labeled with PO1 (Figure 3C).

**Figure 3.**
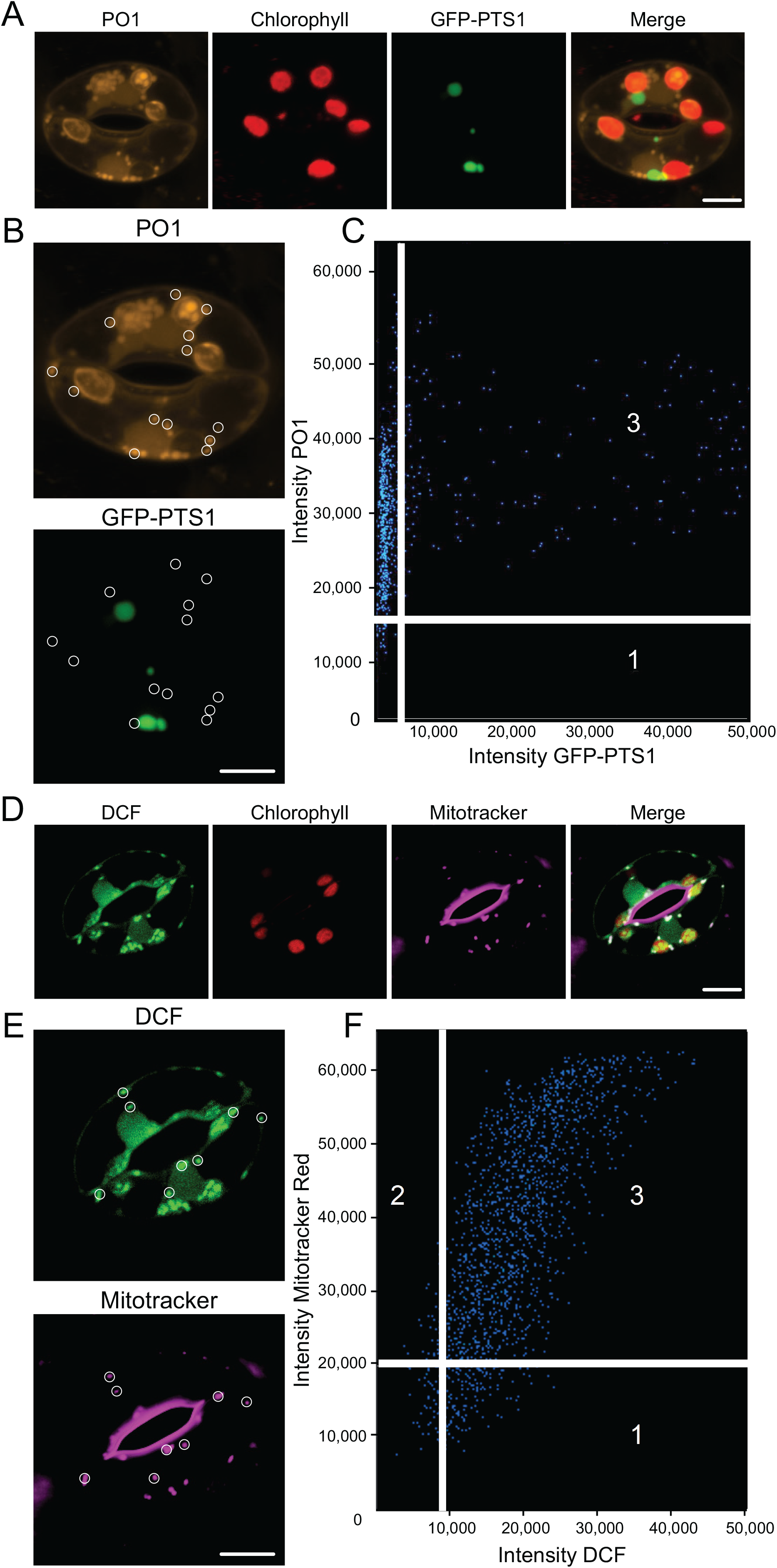
ABA treatment results in increased ROS accumulation in cytosolic puncta that colocalize with mitochondria. A) Confocal micrographs of PO1 fluorescence (orange), chlorophyll autofluorescence (red), GFP-PTS1 (green), and a merged image. Maximum intensity projection of full z-stack is shown. B) Regions of interest used to generate weighted colocalization coefficient are circled in white, highlighting the absence of PO1 fluorescence colocalizing with GFP-PTS1 fluorescence. C) Colocalization graph generated with the ZEN Black colocalization module from regions of interest highlighting PO1-labeled cytosolic puncta. Numbers on scatterplot represent data points that either fall below the determined intensity cutoff for PO1 (1) or GFP-PTS1 (2), or data points that are above thresholding limits for both fluorescent reporters (3). D) Confocal micrographs of DCF fluorescence (green), chlorophyll autofluorescence (red), Mitotracker (magenta), and a merged image showing DCF colocalized with Mitotracker (white). E) Regions of interest used to generate weighted colocalization coefficient are circled in white, showing DCF fluorescence colocalizing with Mitotracker fluorescence. F) Colocalization graph generated with the ZEN Black colocalization module from regions of interest highlighting DCF-labeled cytosolic puncta. Numbers on scatterplot represent data points that either fall below the determined intensity cutoff for Mitotracker (1) or DCF (2), or data points that are above thresholding limits for both fluorescent reporters (3). Scale bars: 5μm.

To determine whether these puncta colocalize with mitochondria, we labeled Col-0 guard cells with Mitotracker Red prior to staining with CM H_2_DCF-DA. Figure 3D shows Mitotracker (magenta), DCF signal (green), and chlorophyll signal (red), separately and in an overlay. All of the DCF puncta contained Mitotracker red signal. We drew ROIs around more than 40 DCF puncta (Figure 3E). Also, these structures display higher DCF intensity than most other localizations which allowed us to use the lowest intensity puncta to define the colocalization threshold and generate a colocalization graph (Figure 3F). A majority of pixels in the designated ROIs contain both DCF and Mitotracker Red signals and we calculated the average weighted colocalization coefficient to be 0.96. These results are consistent with the punctate structures showing ABA-dependent ROS changes being mitochondria.

### The Genetically-Encoded ROS Biosensor roGFP2-Orp1 Showed Rapid ABA-Dependent H_2_O_2_ Increases Within Guard Cell Nuclei and Cytosol

Although chemical ROS probes have provided us with a framework to identify subcellular locations in which ABA drives ROS increases in guard cells, these sensors suffer from disadvantages such as irreversibility and differential dye uptake into some subcellular compartments (Martin et al., 2022). Thus, more precise tools are necessary to reliably evaluate ABA-dependent oxidation in guard cells. Therefore, we evaluated changes in H_2_O_2_ using the genetically encoded biosensor, roGFP2-Orp1, which has enhanced sensitivity relative to PO1, can be targeted to different subcellular locations, and provides a ratiometric readout that is not affected by changes in pH (Nietzel et al., 2019). In the presence of H_2_O_2_, the yeast peroxidase Orp1 protein (also known as glutathione peroxidase 3) becomes oxidized to sulfenic acid (Cys-SOH) on a reactive cysteine residue that rapidly forms an intramolecular disulfide bond with a nearby cysteine. This disulfide is then efficiently transferred via thiol-disulfide exchange to a pair of cysteines on roGFP2, resulting in a conformational change that alters the optical properties of the fluorophore (Gutscher et al., 2009). When reduced, the sensor has increased signal intensity after excitation with the 488 nm laser line, while oxidation leads to elevated signal following 405 nm excitation. Therefore, dividing signal intensity after 405 nm excitation by intensity following 488 nm excitation provides a ratiometric readout which has an internal control for expression levels within a particular tissue (Nietzel et al., 2019).

To identify the dynamic range of this sensor in guard cells, we treated with 20 mM DTT to reduce this biosensor, which leads to a low 405/488 fluorescence ratio (Supplemental Figure S6). In contrast, treatment with 10 mM H_2_O_2_ increases protein oxidation leading to an elevated 405/488 fluorescence ratio. To examine the effect of ABA on oxidation of this sensor, we excised fully mature Arabidopsis leaves containing roGFP2-Orp1 and generated epidermal leaf peels as described above (Figure 4). The process of generating an epidermal leaf peel is a mechanical stress, that can increase roGFP2-Orp1 oxidation (Scuffi et al., 2018). We verified that incubation of the epidermal leaf peels in stomatal opening buffer for 4 h prior to any treatment allowed oxidation to begin returning to baseline levels. Stomatal opening buffer was removed following equilibration and replaced with a similar solution containing 20 μM ABA or a control treatment for 45 min. Figure 4A shows the fluorescence of roGFP2-Orp1 guard cells excited at either 405 or 488 and illustrates that consistent with oxidation of this reporter after ABA treatment, the fluorescence of a sample excited at 405 is elevated and the fluorescence of the sample excited at 488 is reduced. The increase in oxidation after ABA treatment is most evident when the ratio of fluorescence is illustrated as a heat map, with this being generated by an ImageJ plugin (Fricker, 2016) (Figure 4A). We quantified the effect of ABA treatment in roGFP2-Orp1 oxidation in whole stomata, which resulted in a 1.4-fold increase in oxidation ratio when compared to control guard cells (Figure 4B). The dynamic range of the sensor as judged by DTT and H_2_O_2_ treatment defining the minimum and maximum is noted on the graph in Figure 4B-4E. These data are consistent with ABA driving an increase in guard cell H_2_O_2_ during stomatal closure.

**Figure 4.**
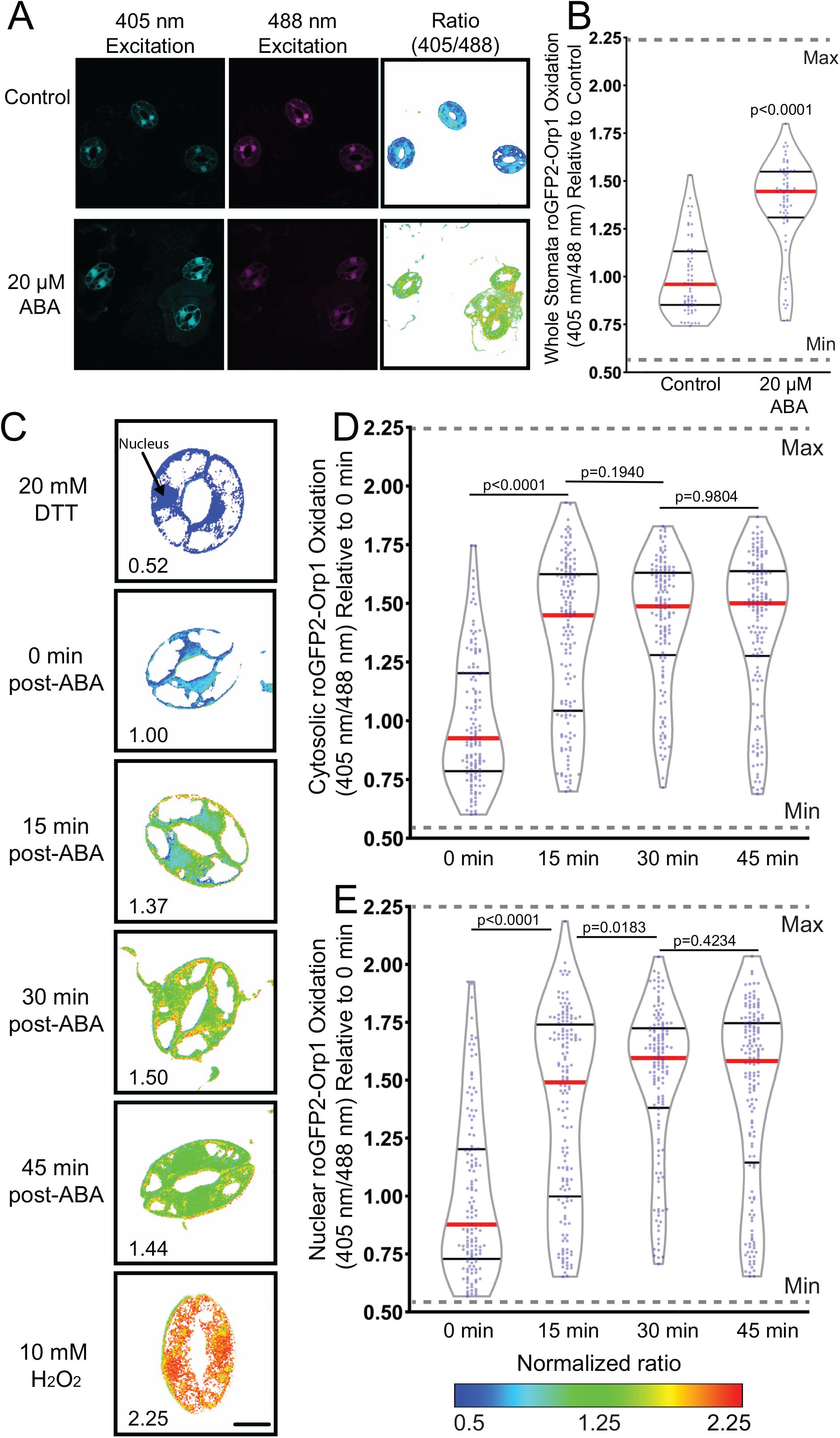
roGFP2-Orp1 detects ABA-increased H_2_O_2_ within the guard cell cytosol and nuclei. A) Confocal micrographs of Arabidopsis guard cells expressing roGFP2-Orp1 treated with 20 μM ABA for 45 min after excitation with either 405 or 488 nm laser line are shown along with ratiometric images that display fluorescence ratios calculated from those images. B) Quantification of intracellular roGFP2-Orp1 ratios following 20 μM ABA or control treatment. Ratios are the fluorescence intensity collected after excitation at 405 nm divided by the intensity after 488 nm excitation. All individual values were normalized to the average of the control treatment and are displayed on the graph as blue dots with the median shown in red and lower and upper quartiles indicated in black with data from three separate experiments (n=64-69) whole stomata for each treatment). P-values were calculated from student t-test. C) Confocal micrographs of Arabidopsis guard cells converted to ratiometric values from cells expressing roGFP2-Orp1 treated with 20 μM ABA for 0, 15, 30, or 45 min. Minimum and maximum sensor oxidation are shown by treatment with 20 mM DTT or 10 mM H_2_O_2_, respectively. Ratios are calculated as above. Normalized ratios are then created relative to the average for the 0 min timepoint. D) Quantification of roGFP2-Orp1 ratio in the cytosol and E) nucleus following 20 μM ABA for 0, 15, 30, or 45 min. Data are reported from three separate experiments (n>131 guard cells for each time point). Minimum and maximum sensor oxidation is represented on graphs by gray dashed lines. The significance of differences between indicated time points were determined by one-way ANOVA followed by a Tukey’s multiple comparisons test and are shown on the graph. Scale bar: 5μm.

A potential benefit of using this genetically encoded sensor is the ability to monitor changes in H_2_O_2_ within individual guard cells over time to gain insights into the spatial dynamics of ABA-dependent increases in H_2_O_2_. The changing sensor oxidation over time in these guard cells treated with ABA and then treated with DTT to reverse this oxidation is shown in Supplemental Figure S7A. The challenge of continuous illumination of the same guard cells is that it can lead to light-induced oxidation of the sensor due to excitation of chloroplasts leading to increases in H_2_O_2_ (Ugalde et al., 2021). Consistent with this prior report, we can initially detect differences in sensor oxidation between ABA treated and buffer control samples, but after 30 minutes of imaging, the amount of oxidation of the sensor in the control became similar to the ABA-induced oxidation (Supplemental Figure S7B). Therefore, rather than time course imaging, we minimized the effect of light on sensor oxidation by imaging multiple different samples at several time points as we did previously with chemical probes.

To examine the temporal dynamics of H_2_O_2_ accumulation within the cytosol and nucleus following ABA treatment, we monitored shifts in oxidation ratio within guard cells transformed with roGFP2-Orp1 by drawing ROIs in these locations. Although this roGFP2-Orp1 protein fusion is targeted to the cytosol, the biosensor also shows signal in the guard cell nucleus (Nietzel et al., 2019; Babbar et al., 2021), allowing us to also monitor H_2_O_2_ changes in this organelle. Leaves expressing roGFP2-Orp1 were peeled and equilibrated in stomatal opening solution as described above. Opening buffer was then removed following equilibration and replaced with 20 μM ABA for 0, 15, 30, or 45 min (Figure 4C). Though roGFP2-Orp1 was still slightly oxidized in both the cytosol and nucleus after 4 hr incubation in stomatal opening solution, treatment with 20 μM ABA resulted in a significant increase in oxidation above baseline in both locations within 15 min (Figure 4D-E). ABA treatment led to continued oxidation of the sensor in a time-dependent manner, reaching a maximum of 1.4-fold increase over control in the cytosol and 1.5-fold in the nucleus at 30 min with the oxidation ratio beginning to decrease at 45 mins (Figure 4C-E).

### ABA Treatment Increases the Oxidation of roGFP2-Orp1 targeted to the Mitochondria and Chloroplast

To confirm that ABA perception drives H_2_O_2_ accumulation in guard cell mitochondria, we examined a transgenic line expressing roGFP2-Orp1 specifically in the mitochondrial matrix (mt-roGFP2-Orp1) (Nietzel et al., 2019) (Figure 5). To minimize oxidation due to generation of the leaf peel, we equilibrated samples in stomatal opening solution for 4 hours before imaging. We determined the dynamic range of this reporter using DTT to fully reduce the reporter and H_2_O_2_ to fully oxidize it, as shown in Supplemental Figure 9 and Figure 5B. It is evident that the signal of this reporter is dispersed in puncta throughout the cytosol, which is most evident in the H_2_O_2_ treated samples. Since the emission spectra of GFP following excitation with its optimal wavelength (488 nm) is easily unmixed from that of PO1 emission at the same excitation wavelength, we utilized this sensor to verify that PO1 labeled puncta were also identified as mitochondria through colocalization of these two signals (Supplemental Figure S8).

**Figure 5.**
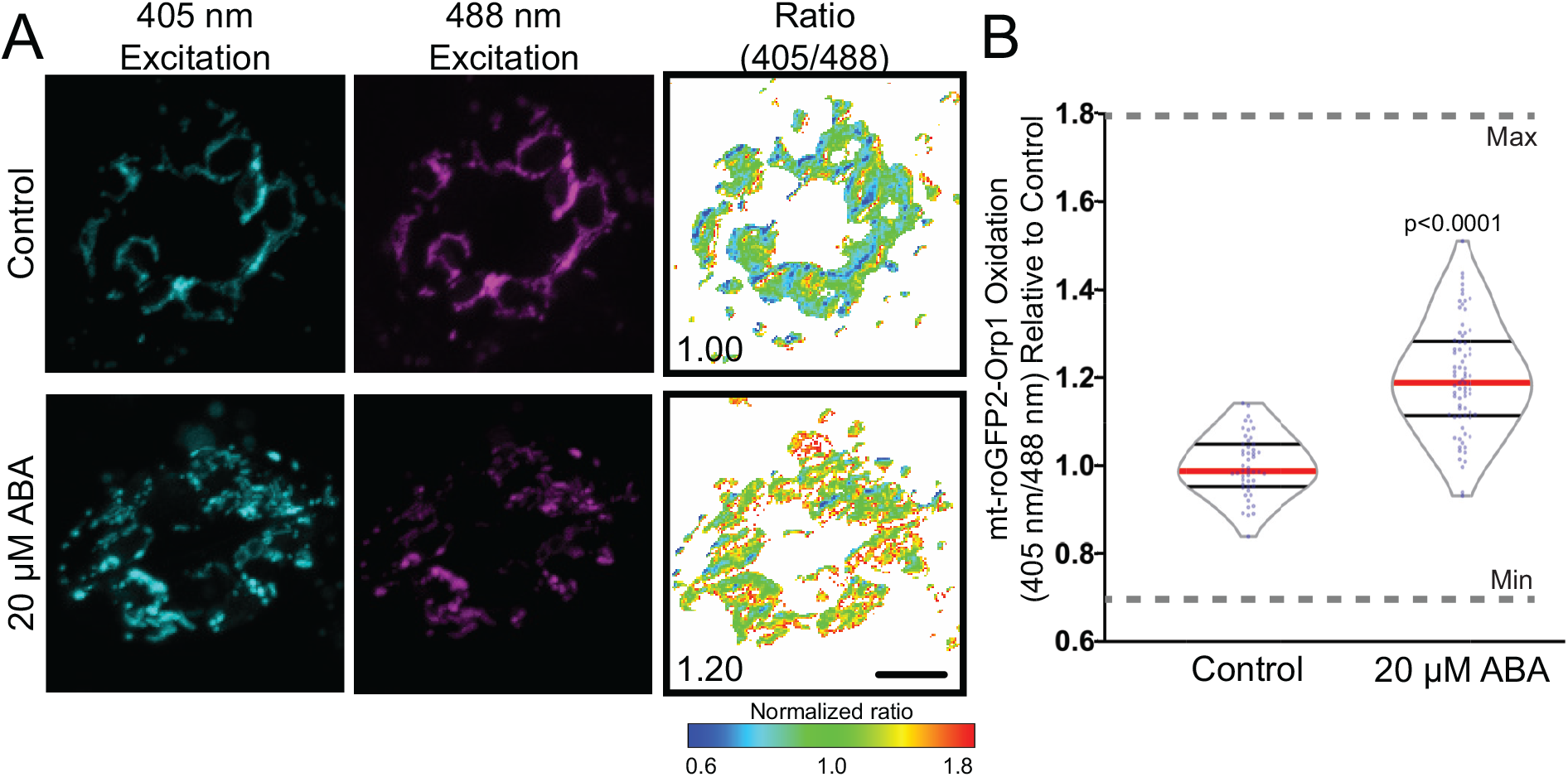
Mitochondrially targeted roGFP2-Orp1 reveals ABA-dependent H_2_O_2_ increases. A) Confocal micrographs of Arabidopsis guard cells expressing mt-roGFP2-Orp1 treated with 20 μM ABA or control buffer for 45 min. Ratiometric images display fluorescence ratios calculated from separate images taken using sequential excitation at 488 nm and 405 nm for each time point. Ratios are calculated by dividing fluorescence intensity collected at emission window 500-535 nm after excitation at 405 nm by the intensity collected in the same emission window after 488 nm excitation. Scale bar: 5μm. B) Quantification of mt-roGFP2-Orp1 ratio in the entire guard cell following 20 μM ABA or buffer control for 45 min. All individual values were normalized to the average of the control treatment and are displayed on the graph as blue dots with the median shown in red and lower and upper quartiles indicated in black. Minimum and maximum sensor oxidation are shown by treatment with 20 mM DTT or 10 mM H_2_O_2_, respectively. Data are reported from three separate experiments (n>50 stomata). Minimum and maximum sensor oxidation, determined by treatment with DTT and H_2_O_2_, respectively, is represented on graphs by gray dashed lines. Listed p-values were determined by one-way ANOVA followed by Tukey’s post hoc test. Scale bars: 5μm.

Treatment with 20 μM ABA resulted in a significant increase in this mitochondrial sensor at 45 minutes after ABA treatment (Figure 5B). It is of note that the mt-roGFP2-Orp1 oxidation state across the entire mitochondrial population is less uniform than with the sensor targeted to the cytoplasm (Figure 5C). This may be responsible for the lower magnitude change in response to ABA in mt-roGFP2-Orp1 as compared to the ABA increase mitochondrial signal of chemical ROS probes.

We also examined the effect of ABA on roGFP2-Orp1 targeted to the chloroplast (plastid-roGFP2-Orp1) (Ugalde et al., 2021). We did detect a significant increase in sensor activity, supporting our result with chemical sensors (Supplemental Figure S10). However, ABA induced a lower magnitude response in this reporter than seen via DCF, suggesting another ROS type or disproportionate localization of the chemical probe to this organelle. The magnitude of the ABA changes in these two sensors in the mitochondria and chloroplasts cannot be directly compared because of multiple technical differences, but in both organelles we see ABA responses mirroring that seen with chemical reporters of ROS changes. Altogether, these results use a sophisticated biosensor to demonstrate that ABA increases H_2_O_2_ with similar spatial and temporal responses to ROS changes detected with chemical sensors, strengthening evidence for ROS as second messengers in ABA-dependent stomatal closure.

### Mutations or Treatments that Alter Mitochondrial ROS Accumulation Influence ABA-Dependent Stomatal Closure

To evaluate the function of ABA-induced ROS in guard cell mitochondria, we searched for mutants with altered ABA response tied to mitochondrial function. The ABA overly sensitive 6 (*abo6)*, was identified in a genetic screen evaluating the ability of ABA to inhibit primary root elongation (He et al., 2012). Consistent with an enhanced response to ABA, this mutant displayed drought tolerance (He et al., 2012). The *abo6* mutation maps to a gene encoding a mitochondrial DEXH box RNA helicase that functions in the splicing of several transcripts that are required for proper function of complex I in the mitochondrial electron transport chain and the protein product of this gene is only expressed in mitochondria (He et al., 2012). Because complex I is a major source of ROS production, impairment at this site can result in increased electron leakage and thus elevated mitochondrial ROS. To ask whether the *abo6* mutant was also enhanced in ABA response in guard cells, we examined the levels of DCF fluorescence in the mitochondria in guard cells of *abo6* (Figure 6A), finding that *abo6* contained 1.3-fold higher levels of DCF fluorescence in guard cell mitochondria when compared to Col-0 under control conditions. Additionally, ABA treatment yielded 1.2-fold higher DCF signal in the *abo6* mutant background than in Col-0 and had a larger ABA response as compared to its untreated control than Col-0 (Figure 6B). These results demonstrate that *abo6* has an enhanced ABA response in guard cells, consistent with the elevated root ABA responses.

**Figure 6.**
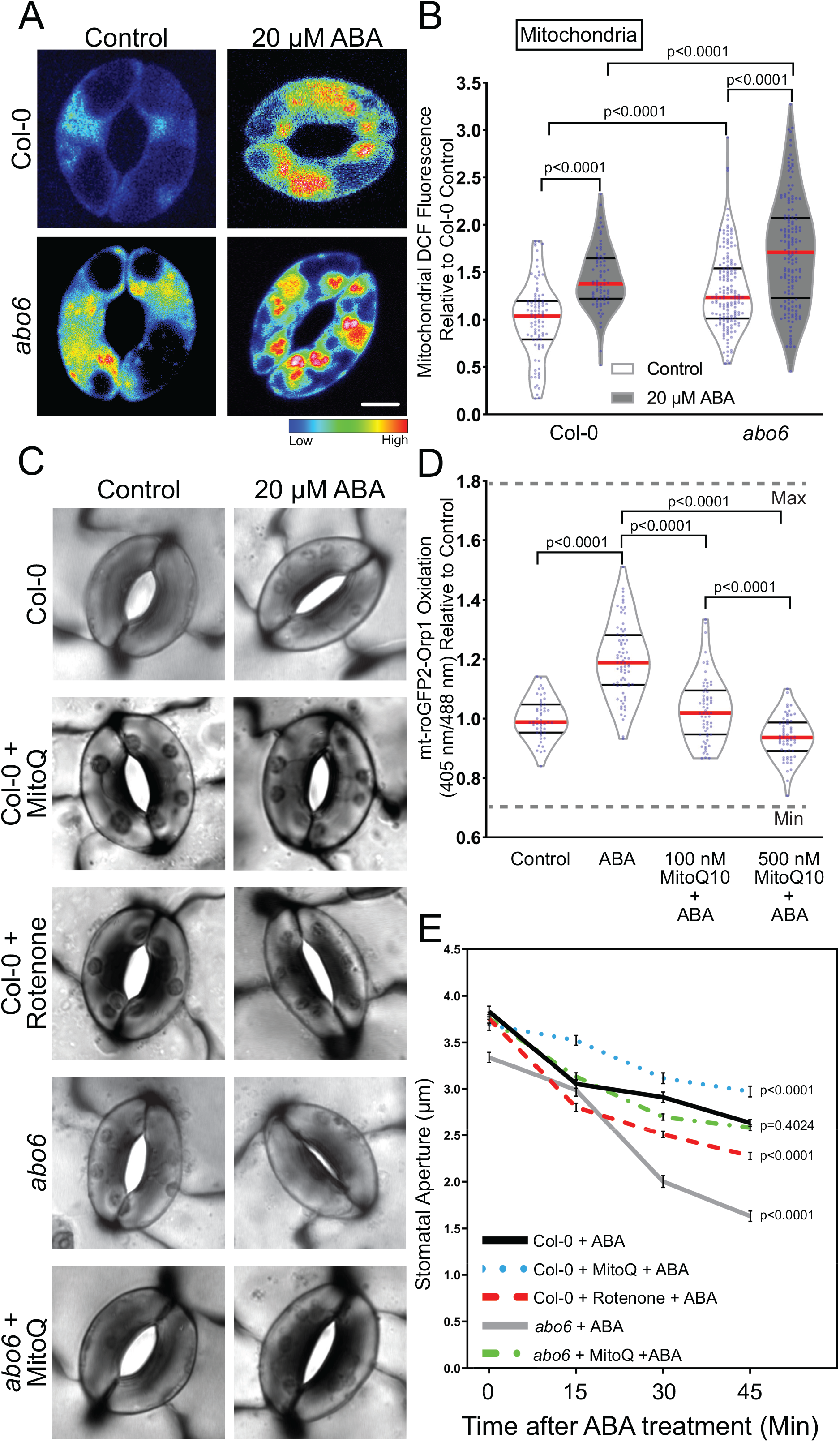
Perturbations in mitochondrial ROS influence the rate of ABA-induced stomatal closure. A) Confocal micrographs of DCF fluorescence following conversion to LUT for *abo6* guard cells treated with control buffer or 20 μM ABA for 45 min. Scale bar: 5μm. B) DCF fluorescence was quantified within mitochondria of Col-0 and *abo6* guard cells with and without ABA treatment from three separate experiments and is reported relative to untreated Col-0, with each bar represented by (n>75) guard cells. C) Stomatal apertures of leaves of Col-0 or *abo6* pretreated with either control buffer, 50 μM rotenone for 1 hr or 500 nM MitoQ for 3 hrs and then treated with 20 μM ABA for 45 min. Scale bar: 5μm. D) Quantification of mt-roGFP2-Orp1 ratio of the entire guard cell following 20 μM ABA, buffer control, or pretreatment with either 100 or 500 nM MitoQ for 3 hrs followed by ABA treatment for 45 min (n=65). E) Stomatal apertures of Col-0 and *abo6* leaves were quantified at 0, 15, 30, 45 min after ABA treatment (n>85 stomata/per reported value) in the presence and absence of MitoQ or rotenone, with the average and SEM graphed at each time point. All individual values in B) and D) were normalized to the average of the control treatment for Col-0 and are displayed on the graph as blue dots with the median shown in red and lower and upper quartiles indicated in black. The p-values for each quantification were generated by two-way ANOVA of the entire time course for each genotype/treatment, followed by Tukey’s post hoc test.

To examine the effect of mitochondrial ROS on guard cell ABA sensitivity, we examined ABA-induced stomatal closure in *abo6* guard cells and in Col-0 guard cells with pharmacological perturbations of mitochondrial ROS (Figure 6C). Here we utilized rotenone, an inhibitor of complex I in the mitochondrial electron transport chain (Palmer et al., 1968) and can result in increased electron leakage out of complex I and ultimately increased accumulation of mitochondrial ROS (Li et al., 2003; Zhou et al., 2014; Mohammed et al., 2020). We also used the mitochondrially targeted antioxidant MitoQ (Kelso et al., 2001), to determine its effect on ABA increased oxidation of mt-roGFP2-Orp1. We treated guard cells with 500 nM MitoQ in advance of ABA treatment. This treatment abolished the ABA-induced increase in mt-roGRP2-Orp1 fluorescence (Figure 6D). We were unable to find reports of this inhibitor being used in plants, suggesting that its specificity toward mitochondria in photosynthetic organisms has not been adequately tested. MitoQ has structural similarity with plastoquinone found in the chloroplast, so we examined MitoQ’s effect on ABA-dependent oxidation of plastid-roGFP2-Orp1. MitoQ lead to a reduction in the oxidation of chloroplast plastid-roGFP2-Orp1 sensor as compared to the ABA only treatment, but not to levels of untreated leaves (Supplemental Figure S11). This MitoQ effect was substantially smaller than the effect on the mitochondrial sensor, which completely reversed the effect of ABA resulting in levels of sensor oxidation that were lower than in the absence of ABA treatment (Figure 6D).

To evaluate the effect of Mito Q on stomatal closure, leaves were excised and peeled and then pretreated with either stomatal opening buffer as described previously, 500 nM MitoQ for 3 hrs, or 50 μM rotenone for 1 hr. Epidermal peels were then treated with 20 μM ABA for 0, 15, 30, or 45 min and guard cells were immediately imaged (Figure 6C). Initial apertures after incubation in opening solution showed a slight difference between Col-0 and *abo6* prior to any ABA treatment, consistent with the elevated levels of baseline mitochondrial ROS observed in Figure 6B. Following 20 μM ABA treatment, *abo6* showed a significant increase in ABA dependent closure relative to Col-0. Unlike *abo6*, rotenone pre-treatment alone did not significantly alter initial stomatal aperture measurements prior to treatment with ABA. However, rotenone pre-treatment significantly increased the amount of stomatal closure over the 45-minute time course of ABA treatment. We also examined the effect of pretreatment with MitoQ on stomatal closure, finding that it significantly reduced ABA-dependent stomatal closure in Col-0 and was able to rescue the hypersensitive ABA response in *abo6* guard cells so that the stomatal aperture was not significantly different from Col-0 (Figure 6E). These findings are consistent with ABA driving ROS increases in guard cell mitochondria and that the degree of ABA sensitivity is correlated with the amount of ROS production within this organelle.

### Mutants Deficient in *rbohd* and *rbohf* Have Impaired ABA-Induced ROS Accumulation in Several Subcellular Locations

RBOH enzymes are well-characterized producers of signaling ROS that regulate a myriad of plant developmental processes and environmental responses (Chapman et al., 2019). To determine whether RBOH enzymes are linked to the increase in ROS accumulation in mitochondria following ABA treatment, we examined DCF fluorescence in Arabidopsis mutants with defects in the genes encoding RBOHD and RBOHF, which were previously reported to function in ABA-induced stomatal closure (Kwak et al., 2003). Figure 7A contains images of DCF fluorescence reported as lookup tables in Col-0 and the double mutant *rbohd/f*. The *rbohd/f* double mutant not only exhibited reduced DCF accumulation in the cytosol (Supplemental Figure 12) following 20 μM ABA treatment for 45 minutes (as predicted by the known function of RBOH proteins in controlling cytosolic ROS), but we also failed to observe a significant increase in signal within the mitochondria of this mutant (Figure 7B).

**Figure 7.**
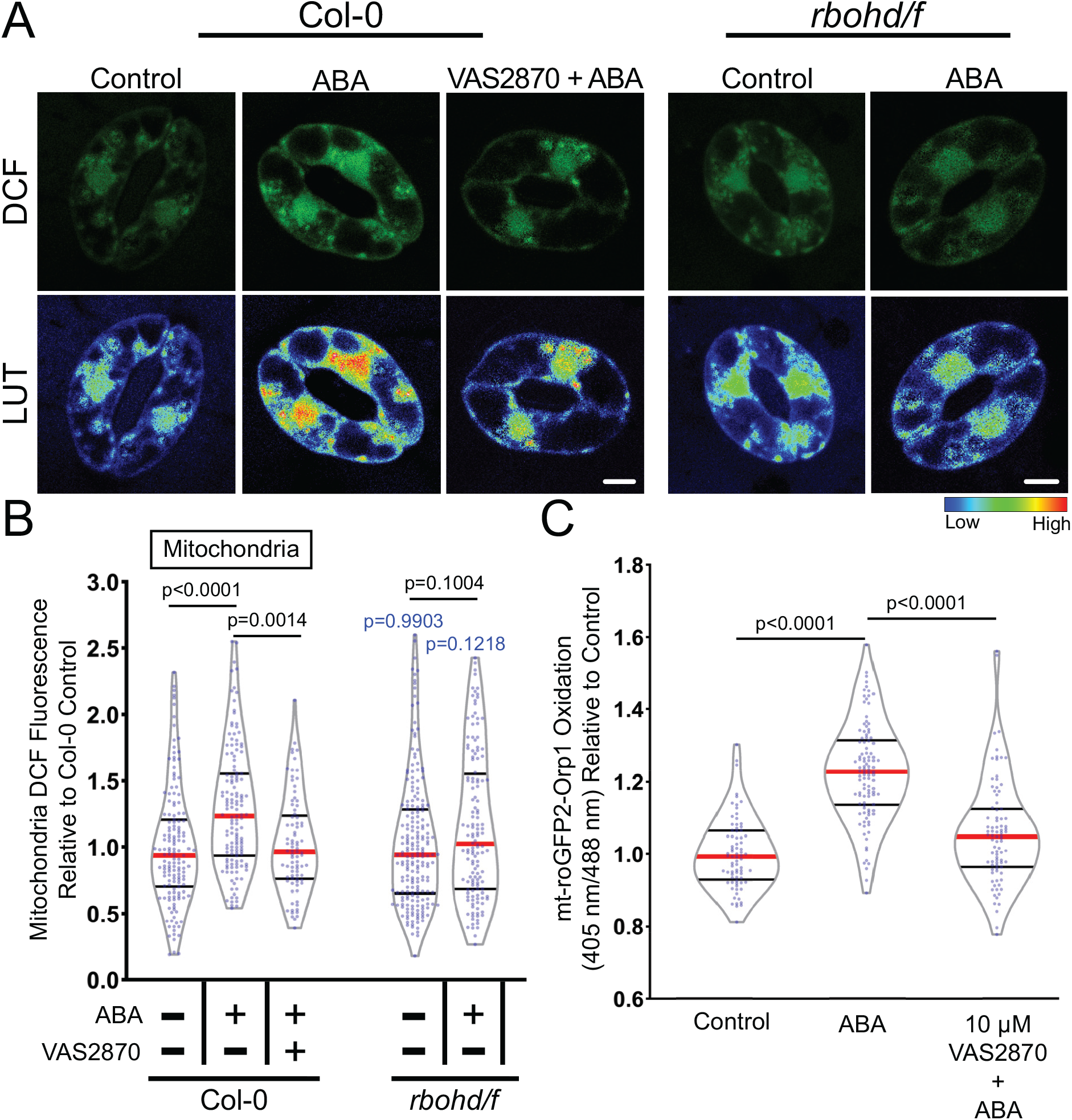
RBOH enzymes contribute to ABA-increased ROS accumulation in guard cell mitochondria. A) Confocal micrographs of DCF fluorescence or DCF images converted to LUT of Col-0 or *rbohd/f* guard cells treated with buffer control or 20 μM ABA as well as Col-0 pre-treated with 10 μM VAS2870 followed by ABA treatment. Scale bar: 5μm. B) Violin plots show quantifications of mitochondrial DCF fluorescence following treatment with control buffer, ABA, or pre-treated with VAS2870 and then treated with ABA from three separate experiments (n>85). C) Quantification of mt-roGFP2-Orp1 ratio of the entire guard cell following 20 μM ABA, buffer control, or pretreatment with 10 μM VAS2870 followed by ABA treatment (n>77). All individual values were normalized to the average of the control treatment in Col-0 and are displayed on the graph as blue dots with the median shown in red and lower and upper quartiles indicated in black. P-values in black font represent the significance of differences between treatments in the same genotype spanning the compared treatments. P-values in blue font representing the significance of differences between *rbohd/f* and Col-0 under the same treatment conditions. P-values are recorded according to two-way ANOVA followed by Tukey’s post hoc test.

We also inhibited RBOH-dependent ROS production with a pharmacological approach. Although diphenylene iodonium (DPI) is a commonly used NADPH-oxidase inhibitor that inhibits ABA dependent guard cell closure in guard cells (Zhang et al., 2001; Gayatri et al., 2017; Watkins et al., 2017), the molecule is a general flavoprotein inhibitor that can directly interfere with metabolic processes in mitochondria (Augsburger et al., 2019). Therefore, we utilized the more selective pan NOX inhibitor VAS2870, which selectively targets an active-site cysteine that is conserved in mammalian NOX enzymes and plant RBOHs (Yun et al., 2011). We confirmed that VAS2870 regulated ABA responses in guard cells by evaluating its effect on ABA-induced stomatal closure. Pre-treatment with 10 μM VAS2870 for 1 hr prior to ABA treatment was able to significantly inhibit ABA-dependent stomatal closure compared to guard cells treated with ABA alone, suggesting it is an effective RBOH inhibitor in guard cells (Supplemental Figure S12A-B) and supporting the requirement of RBOH activity for full ABA-induced stomatal closure. Pre-incubation with VAS2870 abolished ABA-induced ROS increases in both the mitochondria (Figure 7B) and the cytosol (Supplemental Figure 12). We also pretreated mt-roGFP2-Orp1 guard cells with VAS2870 to verify that the ABA-dependent increases in mitochondrial oxidation seen previously were diminished by inhibitor treatment (Figure 7C). Altogether, these findings indicate that RBOH enzymes play a role in ABA-dependent ROS accumulation in guard cell mitochondria.

## Discussion

Plants regulate stomatal aperture in response to environmental and hormonal signals through the control of guard cell turgor pressure (Nilson and Assmann, 2007). During drought stress, plants increase ABA synthesis (Muhammad Aslam et al., 2022), which initiates a complex signaling pathway that ultimately leads to stomatal closure (Vishwakarma et al., 2017). Included in this signaling cascade is the activation of RBOH enzymes that trigger a burst of ROS that act as second messengers in guard cell closure (Kwak et al., 2003). The accumulation of ROS in guard cells following ABA treatment has been detected primarily by increases in DCF fluorescence, a nonspecific chemical ROS sensor, with signal quantified across the whole guard cell (Pei et al., 2000; Watkins et al., 2014; Watkins et al., 2017; Postiglione and Muday, 2020). However, signaling through ROS increasingly appears to be much more nuanced, with signaling outputs depending on both which ROS is generated and where it is generated (Noctor and Foyer, 2016). Recent technological advances in microscopic resolution and ROS detection (Winterbourn, 2014; Nietzel et al., 2019; Ugalde et al., 2021), as well as a growing number of genetic resources to disrupt signaling and/or ROS synthesis allowed us to ask more precise questions about the functions of ROS, including when and where these are made and which ROS function to control signaling and development (Martin et al., 2022). In this study, we use both ROS-responsive fluorescent dyes and genetically encoded biosensors to demonstrate that ABA increases H_2_O_2_ within specific subcellular compartments of guard cells and that these changes are both necessary and sufficient to drive stomatal closure.

We found that ABA significantly increased the fluorescence intensity of the generic ROS probe, DCF, in the cytoplasm, mitochondria, nuclei, and chloroplasts, with the most substantial increases in the mitochondria and chloroplast. The identity of these organelles was verified by linear unmixing DCF signal from organelle localized dyes and chloroplast autofluorescence. The ABA-induced ROS synthesis in the mitochondria, cytosol, and nucleus were all dependent on the canonical ABA signaling pathway, as the ABA-increased ROS was lost in these organelles in the quintuple ABA receptor mutant *pyl-11458*. However, ABA was still able to trigger a significant increase in the chloroplasts of *pyl-11458* suggesting alternate mechanisms for ROS generation in this organelle during ABA signaling.

A major component of these ABA-induced ROS is due to increases in H_2_O_2_. We used the H_2_O_2_-selective chemical probe, PO1, and the H_2_O_2_ responsive genetically-encoded sensor roGFP2-Orp1, targeted to the cytosol and nucleus, to the mitochondria, or to the chloroplast. This highly reactive biosensor contains a ratiometric readout which provides advantages over chemical dyes due to its self-normalizing output (Gutscher et al., 2009; Nietzel et al., 2019; Ugalde et al., 2021). This sensor has a cysteine residue that is oxidized by H_2_O_2_ leading to a conformational change that alters its optical properties increasing its signal after excitation at 405 nm. Using this sensor, we reveal that ABA treatment leads to a significant increase in H_2_O_2_ when oxidation ratio in the whole guard cell, cytosol, nucleus, are quantified or when mitochondrial or chloroplast ROS are quantified using the targeted mt-roGFP-Orp1 or plastid-roGFP-Orp1. We also detected ABA-dependent increases in H_2_O_2_ in the chloroplast and in mitochondria with PO1, but not in the cytosol. We stained leaves containing the mitochondrially targeted roGFP2-Orp1 with PO1 and confirmed that the mitochondria labeled with PO1 displayed strong colocalization with mt-roGFP2-Orp1. Together these experiments reveal ABA-dependent H_2_O_2_ increases in the mitochondria and chloroplasts.

We used several approaches to ask if mitochondrial ROS increases are able to facilitate ABA dependent stomatal closure. A previous study screened mutants for altered ABA-dependent primary root growth (He et al., 2012; Yang et al., 2014). This approach identified the *abo6* mutant, which was isolated for enhanced ABA response in root cells and had drought tolerance, and elevated ROS within root cells. The mutant gene encoded a DEXH box RNA helicase, which led to altered synthesis of mitochondrial electron transport proteins and the gene product is localized to the mitochondria of leaf protoplasts (He et al., 2012). The *abo6* mutant had elevated mitochondrial ROS in guard cells with and without ABA treatment and significantly enhanced ABA-induced stomatal closure relative to Col-0. We also observed enhanced ABA-sensitivity in Col-0 guard cells pretreated with mitochondrial complex I inhibitor, rotenone, which also elevates mitochondrial ROS accumulation (Li et al., 2003; Zhou et al., 2014; Mohammed et al., 2020). Rotenone’s mechanism of action made this a particularly intriguing finding as the inhibitor increases mitochondrial ROS while limiting oxidative phosphorylation and ATP synthesis (Palmer et al., 1968). This suggests that our findings of increased ABA sensitivity were largely based on the pool of ROS generated in this organelle and not increased energetic requirements.

We also showed that pretreatment with the mitochondrially targeted ROS scavenger, MitoQ (Kelso et al., 2001), significantly blunted mt-roGFP2-Orp1 oxidation and ABA-induced stomatal closure in Col-0 guard cells. As we found no other reference to MitoQ use in plants, we also asked whether it acted on chloroplasts. Although this compound reduced oxidation of the plastid-roGFP2-Orp1, the effect was smaller than on mt-roGFP2-Orp1. In particular, MitoQ reduced mt-roGFP2-Orp1 levels to below those of ABA treatment, while plastid-roGFP2-Orp1 was still significantly oxidized by ABA even after MitoQ pretreatment, suggesting MitoQ has mitochondrial selectivity.

Additionally, a previous reports demonstrated that impairment of chloroplastic ROS generation through chemical inhibition of photosynthetic electron transport does not have an effect on ABA-induced stomatal closure (Wang et al., 2016). Another study showed that a mutant that lacked chlorophyll in guard cells was still able to close stomata following ABA treatment (Azoulay-Shemer et al., 2015). ABO6 is localized to mitochondria and absent in chloroplast (He et al., 2012) and MitoQ is able to reverse the enhanced ABA stomatal closure phenotype of *abo6* to wild-type levels. Together, these results are consistent with mitochondria as necessary sites of ROS generation for productive ABA signaling in guard cells.

RBOH enzymes were previously implicated in ABA-dependent increases in DCF fluorescence (Kwak et al., 2003; Drerup et al., 2013; Hsu et al., 2018; Postiglione and Muday, 2020) by examination of an *rbohD/rbohF* double mutant (Kwak et al., 2003) and treatment with a nonselective RBOH inhibitor, Diphenyleneiodonium (DPI) (Watkins et al., 2017). RBOH enzymes have well established roles in hormone induced ROS synthesis in mammals (Vermot et al., 2021) and plants (Chapman et al., 2019). These enzymes produce superoxide in the apoplast, which can be rapidly dismutated into H_2_O_2_ via superoxide dismutase. The influx of H_2_O_2_ into the cytoplasm is facilitated by aquaporins, making it available to reversibly oxidize cytoplasmic protein targets to change their conformation, activity and/or regulatory properties (Tian et al., 2016; Rodrigues et al., 2017). Yet, whether RBOH enzymes drive ROS accumulation in guard cells in regions beyond the cytoplasm has not been examined.

In guard cells of both an *rbohd/f* double mutant and cells treated with a highly specific pan-RBOH/NOX inhibitor,VAS2870 (Reis et al., 2020), there were reductions in both ABA-induced cytosolic and mitochondrial ROS and ABA-dependent guard cell closure. This suggests a link between RBOH-dependent ROS production and ROS accumulation in guard cell mitochondria. This finding is consistent with a previous report which found that introduction of the *rbohf* mutation into *abo6* relieved ABA hypersensitivity in *abo6* roots (He et al., 2012). This finding both emphasizes the role of RBOH in ABA-induced ROS in cellular locations beyond the cytosol, such as mitochondria, which is an important insight into the function of this class of signaling driven, ROS synthesizing enzymes. Previous work in mammalian systems has shown the ability of NOX enzymes, such as NOX4, to localize to mitochondria (Dikalov, 2011; Shanmugasundaram et al., 2017). However, whether there are RBOHs localized to mitochondria is not currently known in plants.

The regulation of stomatal aperture in response to environmental stimuli, such as drought, is a crucial process in plant adaptation to stress. This study built on prior evidence that ABA drives ROS increases, identified here as H_2_O_2_ as central to this response, as well as revealing the spatial regulation of these ROS. The addition of genetically encoded biosensors to our toolkit of ROS responsive chemical probes allowed us to overlay the position of ROS accumulation with organelle specific markers to reveal ABA-elevated ROS in the mitochondria and chloroplast. In support of a function of mitochondrially derived ROS in guard cells, in both a mutant with increased mitochondrial ROS and chemically-increased mitochondrial ROS production displayed an increased rate of ABA-induced stomatal closure. Meanwhile, a ROS scavenger targeted to this organelle reduced guard cell ABA sensitivity, suggesting ROS production in this organelle functions in the ABA signaling pathway. We also demonstrated that RBOH enzymes play a role in ABA-increased ROS accumulation not only in the cytoplasm, but also guard cell mitochondria. Together these results indicate that ABA-induced H_2_O_2_ accumulation exhibits tight compartmentalization in organelles such as mitochondria that influences guard cell signaling and physiology.

## Methods and Materials

### Plant Growth Conditions

*Arabidopsis thaliana* seeds that were used include Col-0, *rbohd/rbohf* double mutant (Miller et al., 2009), *pyl1-1*; *pyr1-1*; *pyl4-1*; *pyl5*; *pyl8-1* quintuple mutant *(pyl-11458)* (Zhang et al., 2020) (ABRC), *ABA overly sensitive 6* (*abo6)* (Alonso et al., 2003) (ABRC), GFP-PTS1 reporter (Ramón and Bartel, 2010), roGFP2-Orp1 (Nietzel et al., 2019), and mt-roGFP2-Orp1 (Nietzel et al., 2019). Arabidopsis plants were germinated on 1× Murashige and Skoog medium, pH 5.6, Murashige and Skoog vitamins, and 0.8% (w/v) agar, buffered with 0.05% (w/v) MES and supplemented with 1% (w/v) sucrose. After vernalization at 4°C for 48 h, plates were placed under 24-h 120 *μ*mol m^−2^ s^−2^ cool-white light. Seven days after germination, seedlings were transferred to SunGro Redi-Earth Seed Mix. Plants are then grown under a short-day light cycle (8 h light/ 16 h dark) of 120 *μ*mol m^−2^ s^−2^ cool-white light with relative humidity kept between 60-70%. Experiments were conducted on leaves from plants 3 to 4 weeks after germination, unless noted otherwise.

### DCF Staining, Imaging, and Quantification

CM 2,7-dihydrodichlorofluorescein diacetate (CM-H_2_DCF-DA) was dissolved in dimethyl sulfoxide to yield a 50-μM stock. This was diluted in deionized water to yield a final concentration of 4.3 μM with 0.1% (v/v) dimethyl sulfoxide. Epidermal peels of Col-0, *abo6*, or *rbohd/f* were prepared by evenly spraying a microscope slide with a silicone-based medical adhesive (Hollister stock #7730). After 10 min, the abaxial epidermis of the leaf was pressed into the dried adhesive coat, a drop of water was placed on the leaf surface, and the leaf was gently scraped with a razor blade until only the fixed epidermis remains. Fresh epidermal peels were then fully covered in opening solution (50 mM KCl, and 10 mM MES buffer, pH 6.15) and incubated under cool-white light (120 μmol m^−2^ s^−1^) for 3 hrs. For VAS2870 treatments, leaf opening buffer was then replaced with stomatal opening solution containing 10 μM VAS2870 for 1 hr during the opening process. Epidermal peels were then treated with fresh stomatal opening buffer (control buffer) or a similar solution containing 20 μM ABA for 45 min. Pre-treatments were fully removed, and the epidermis was stained for 15 min with 4.3 μM H_2_DCF-DA stain and washed with deionized water. Microscopy was performed on the Zeiss LSM880 laser scanning confocal microscope with a 32-detector GaAsP array for spectral unmixing. The Plan Apochromat 63x/1.2NA water objective was used for acquisition. The 488 nm laser line was used to excite the leaf surfaces with 0.4% maximum laser power with a 3.5 digital gain. The gain settings were optimized to produce maximum DCF signal while preventing oversaturation. All micrographs were acquired using identical offset, gain, and pinhole settings using the same detectors for each experiment. Settings were defined to spectrally separate the DCF and chlorophyll signal by capturing the emission spectrum for each compound in regions which there was no overlap. Total emission was collected using lambda scanning with a 1 Airy Unit pinhole aperture yielding a 0.9 μm section, the DCF signal alone would later be unmixed from the image for quantification.

Images used for quantification were taken with averaging of 2 with minimal pixel dwell time making sure to limit excess exposure to the laser that may induce ROS. Maximum intensity projections were produced from Z-stacks. The average intensity values within each ROI were acquired and all values obtained were normalized to the average of each subcellular location under control conditions from three biological replicates with 2-3 technical replicates per experiment. DCF fluorescence intensities were measured in ImageJ by drawing ROIs around the whole stomata, chloroplasts, cytosol, nuclei, and individual mitochondria of multiple guard cells.

The images shown in the figures were captured at high resolution using separate but identically treated samples to those used in the quantification with increased averaging, digital zoom, and pixel dwell time to increase resolution and were not included in any quantification. Individual images were selected that were representative of the magnitude of responses in the images generated for quantification. To produce heat maps, we converted pixel intensities of DCF fluorescence using the look-up tables (LUT) function in the Zen Blue Software.

### PO1 Staining, Imaging, and Quantification

Peroxy Orange 1 (PO1) is an H_2_O_2_ sensor, which was dissolved in dimethyl sulfoxide to yield a 5 mM stock. This was diluted in deionized water to yield a final concentration of 50 μM. Epidermal peels were prepared, and guard cells were fully opened as described above, then treated with 20 μM or 100 μM ABA or a control buffer. Pre-treatments were fully removed, and the epidermis was stained for 30 min with 50 μM PO1 dye and washed with deionized water. Microscopy was performed via a Zeiss LSM880 laser scanning confocal microscope with a 32-detector GaAsP array for spectral unmixing. The Plan Apochromat 63x/1.2NA water objective was used for acquisition. Leaf surfaces were excited with the 488 nm laser line at 0.6% maximum laser power and a digital gain set to 3.5. The gain settings were optimized to produce maximum PO1 signal while preventing oversaturation. All micrographs were acquired using identical offset, gain, and pinhole settings using the same detectors for each experiment. Settings were defined to spectrally separate the PO1 signal from chlorophyll autofluorescence by capturing the emission spectrum for each compound in regions where only one signal was present. Total emission was then collected using lambda scanning with a 1 Airy Unit pinhole aperture yielding a 0.9 μm section, the spectral signature that was previously calculated as PO1 signal alone was later unmixed from the image for quantification.

Three-dimensional images of PO1 were acquired on the Zeiss LSM880 system equipped with 32-detector GaAsP array for Airyscan acquisition. Samples were excited with an argon 488 nm laser line using 6% laser power and a Plan Apochromat 63x/1.2NA water objective was used for image acquisition. Total emission was collected using Airyscan of a z-stack spanning the entire depth of a whole guard cell pair, using the optimal optical slice size calculated by the ZEN Black acquisition software. Images were then rendered in three dimensions using Aivia image analysis software. x,y and z,y projections were then acquired of cropped regions containing chloroplasts or mitochondria.

To produce heat maps displayed in Figure 2, we converted pixel intensities of PO1 fluorescence using the look-up tables (LUT) function in the Zen Blue Software. PO1 fluorescence intensities were measured in ImageJ by drawing ROIs around the chloroplasts, cytosol, nuclei, and individual mitochondria of each guard cell. Due to there being more puncta with bright PO1 fluorescence evident after ABA treatment, as reported previously in tomato (Watkins et al., 2017), we obtained a greater number of data points as ABA concentrations increased. The average intensity values within each ROI were acquired and all values obtained were normalized to the average of each subcellular location under control conditions from three biological replicates with three technical replicates per experiment.

### Colocalization analysis of ROS Chemical Probes with Mitochondria and Peroxisomes

For evaluation of peroxisomal colocalization, Arabidopsis leaves containing PTS1-GFP were peeled and labeled with 50 μM PO1 as described above. For evaluation of mitochondrial colocalization, Col-0 Arabidopsis leaves were peeled and treated with 8.6 μM CM H_2_DCF-DA as described above and then 1 μM Mitotracker Red for 15 min. Leaves were then visualized using the Zeiss 880 LSCM device as described earlier. Each signal was resolved in lambda scanning mode, with emission spectra for each individual spectra being obtained prior to colocalization analysis by imaging single labeled samples. Images were taken at multiple Z-positions, though not combined into maximum intensity projections as to not misrepresent colocalization of signals that might be found in the same vertical plane but at different depths. Emission spectra for each signal were then unmixed from corresponding images to better evaluate how either signal contributed to a particular location. For colocalization analysis, samples were examined using the Zeiss Zen colocalization module. The threshold for PO1, GFP-PTS1, DCF, Mitotracker Red, mt-roGFP2-Orp1 was determined in each image, via regions that contained only one fluorescent signal. Regions of interest surrounding mitochondria in PO1 or DCF were then selected and evaluated for colocalization with either GFP-PTS1, mt-roGFP2-Orp1, or Mitotracker Red. Colocalization was then calculated using Pearson’s coefficients (weighted colocalization coefficients) and respective scatterplots were generated.

### Imaging and Analysis of ROS-Sensitive Genetically Encoded Biosensors

Fully mature Arabidopsis rosettes containing roGFP2-Orp1, plastid-roGFP2-Orp1, or mt-roGFP2-Orp1 were excised and peeled prior to being submerged in stomatal opening buffer to equilibrate for 4 hrs to establish a baseline. For inhibitor treatments, leaf peels were pretreated with stomatal opening buffer containing either 500 nM mitoquinol mesylate (MitoQ) for 3 hrs, or 10 μM VAS2870 for 1 hr during the equilibration process. Inhibitor solutions were then removed, rinsed, and replaced with fresh stomatal opening buffer and allowed to incubate until the 4 hr opening period was completed. Stomatal opening buffer or inhibitors were then removed following equilibration and replaced with a similar solution containing 20 μM ABA for 0, 15, 30, or 45 min. Microscopy was performed on the Zeiss LSM880 laser scanning confocal microscope with a Plan Apochromat 63x/1.2NA water objective was used to sequentially excite leaf surfaces at 405 and 488 nm with 1% maximum laser power. Emission was recorded between 505–535 nm to keep autofluorescence at a minimum, with a 2.4 Airy Unit pinhole aperture yielding a 2.0 μm section. We verified that there were no changes in the ratiometric signal calculated with ABA treatment of Col-0 that was not transformed with this biosensor. Images used for quantification were taken without averaging and with scan speed and pixel dimensions optimized for minimal pixel dwell time in order to limit laser-induced oxidation of the sensor. Z-slice number was held constant for all image stacks to promote equity of light exposure (and equal photooxidation) across samples. Z-axis profiles of averaged intensity within a 3 μm^2^ spot size were plotted in ImageJ to verify that both fluorescent channels showed alignment of peak intensity values at proximal stack depths. This check was critical to assure depth alignment of our two distinct fluorescent channels given that we utilized maximum intensity projections for these analyses. Dynamic range of each dye sensor was defined by treating equilibrated samples with 20 mM DTT or 10 mM H_2_O_2_ to determine the maximum reduction or maximum oxidation, respectively.

Images of roGFP2-Orp1 targeted to the cytosol or plastids were captured as described above and maximum intensity projections were analyzed in ImageJ by drawing a region of interest in the nucleus, a cytosolic region, or individual chloroplasts within each guard cell. Images of mt-roGFP2-Orp1 were analyzed by drawing a region of interest around the entire stomata and thresholding to exclude pixels of background intensity values from each measurement. Ratios were calculated by dividing fluorescence intensity following excitation at 405 nm by fluorescence intensity collected after 488 nm excitation. All individual values obtained were normalized to the average of buffer control treated stomata or the 0-minute timepoint in the case of time courses. Ratiometric micrographs were generated using the Ratio Redox Analysis MatLab program package (Fricker, 2016).

### Stomatal Closure Assay

ABA-induced stomatal closure assays were performed with plants 3 to 4 weeks after germination. Epidermal strips were prepared and fully covered in opening solution as described above. For inhibitor treatments, opening buffer was replaced with stomatal opening solution containing either 50 μM rotenone for 1 hr, 500 nM mitoquinol mesylate (MitoQ) for 3 hrs, or 10 μM VAS2870 for 1 hr during the opening process. To induce stomatal closure, opening buffer was replaced with equal volume of a similar solution with 20 μM added to induce closure. For quantification of stomatal aperture, leaf peels were imaged on an ECHO Revolve microscope using transmitted light. Images were acquired using an Olympus UPlanSApochromat 40x/0.95NA objective.

### Statistical Analysis

The data for the subcellular localization of DCF and whole stomata roGFP2-Orp1 quantifications were analyzed via student t-tests using GraphPad Prism 9 comparing ABA to control in each respective cellular location. DCF fluorescence intensities for ABA signaling mutants and widths of stomatal apertures were analyzed by two-way ANOVA, while subcellular PO1 signal intensity, and roGFP2-Orp1 time course data were analyzed by one-way ANOVA using GraphPad Prism 9. This analysis evaluated differences within genotypes between different treatments and compared between genotypes under similar treatments for ABA signaling mutants, between control and multiple ABA treatments for PO1 fluorescence, and between timepoints for roGFP2-Orp1 and mt-roGFP2-Orp1. Tukey’s multiple comparison tests were then utilized to resolve significant differences between treatments, genotypes, or timepoints.

### Accession Numbers

Sequence data from this article can be found in the GenBank/EMBL data libraries under the accession numbers: *PYR1* (At4g17870), *PYL1* (At5g46790), *PYL4* (At2g38310), *PYL5* (At5g05440), *PYL8* (At5g53160), *ABO6* (At5g04895), *RBOHD* (At5g47910), *RBOHF* (At1g64060).

## Acknowledgments

We would like to acknowledge the assistance of Dr Heather Brown Harding (Microscopy core facility) with imaging. We thank Dr. Leslie Poole for generous sharing of expertise and research material. We also thank Antipodean Pharmaceuticals, Inc for supplying mitoquinol mesylate (MitoQ). We greatly appreciate the generosity of Dr Andreas Meyer and Dr Jose Ugalde in sharing roGFP2-Orp1, plastid-roGFP2-Orp1, and mt-roGFP2-Orp1 seeds, as well as providing helpful feedback on an early draft of the manuscript. We also thank Dr Gad Miller for providing *rbohd/f* seeds and Dr. Bonnie Bartel for the GFP-PTS1 transgenic line, and the ABRC for distributing *pyl-11458* and *abo6* seeds. Lastly, we would like to thank members of the Muday lab for valuable input on the manuscript. This work was funded by NSF IOS-1558046 to G.K.M and a fellowship from the Center for Molecular Signaling and an NIH T32 GM127261 fellowship to A.E.P. The content is solely the responsibility of the authors and does not necessarily represent the official view of the National Institutes of Health.

## Author Contributions

AEP designed experiments, performed research, analyzed data, and wrote the paper. GKM designed experiments, wrote, and edited the paper.

