## Supplemental Figures for "Abscisic Acid Increases Hydrogen Peroxide in Mitochondria to Facilitate Stomatal Closure"

### Postiglione and Muday Supplemental Figures

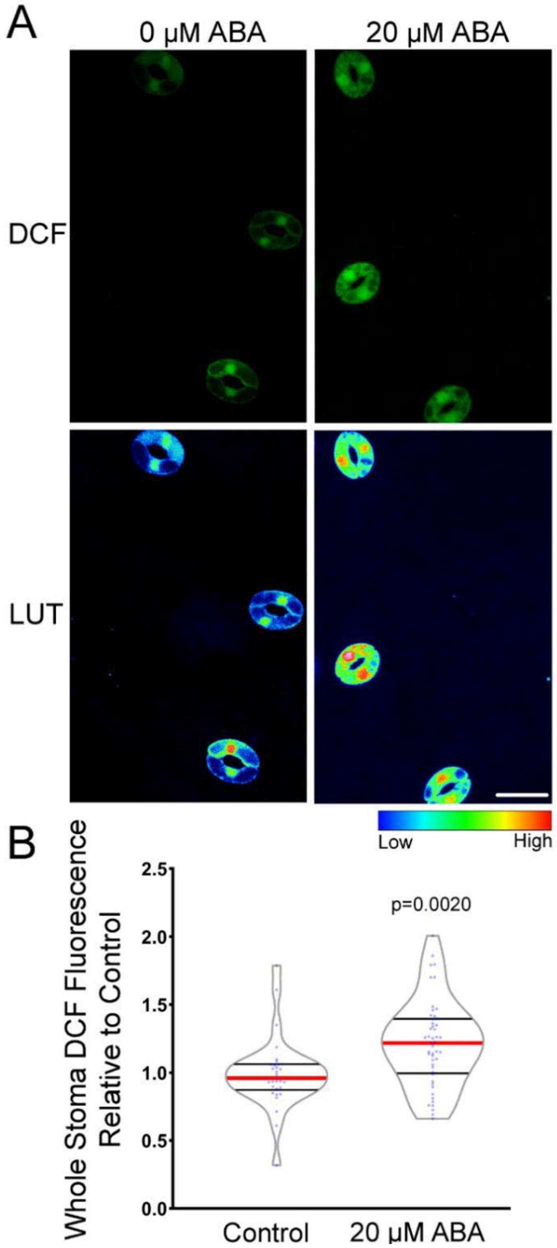


**Supplemental Figure 1.** ABA increases DCF fluorescence when entire Arabidopsis guard cells are quantified.

A) Confocal micrographs of DCF fluorescence of Arabidopsis guard cells treated with buffer control or 20 µM ABA for 45 min shown directly or after conversion to lookup tables (LUT). Scale bar: 20µm. B) Violin plot showing quantifications of DCF fluorescence in the entire guard cell with and without ABA treatment in 30-50 whole stomata per treatment from three separate experiments. All individual values were normalized to the average of the control treatment and are displayed on the graph as blue dots with the median shown in red and lower and upper quartiles indicated in black. P-values from paired student t-tests for samples compared to untreated controls are reported.

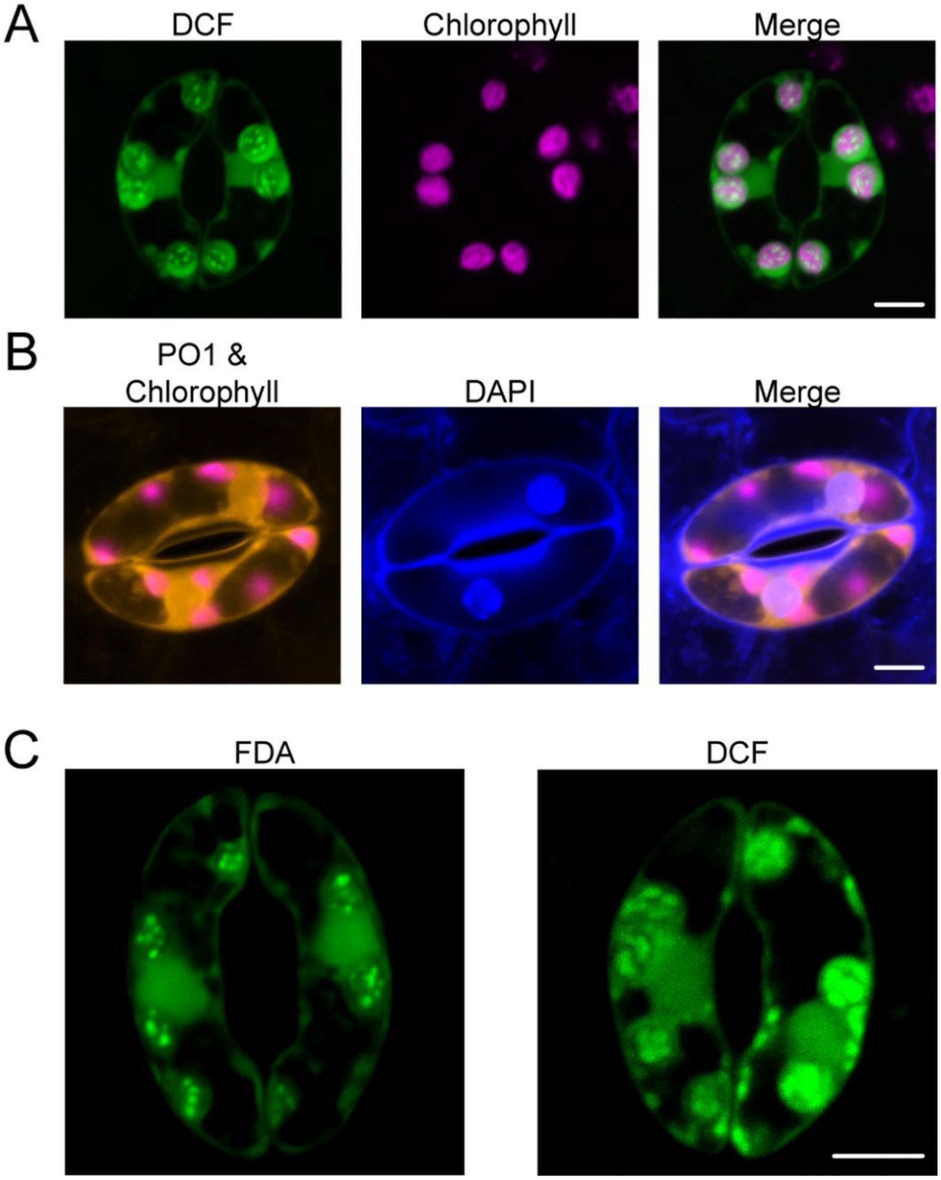


**Supplemental Figure 2.** ROS-sensitive fluorescent probes accumulate in chloroplasts and nuclei.

A) Confocal micrographs of DCF fluorescence (green), chlorophyll autofluorescence (magenta) and merge of both images. B) Confocal micrographs of PO1 (orange) combined with chlorophyll autofluorescence (magenta), DAPI nuclear fluorescence (blue), and a merge of both images. C) Confocal micrograph of Fluorescein Diacetate (FDA), which becomes fluorescent upon cellular uptake, but is not oxidized by ROS and displays similar accumulation pattern as the redox-sensitive variant of the probe Dichlorofluorescein (DCF). Maximum intensity projections of full z-stacks are shown in for each image. Scale bar: 5μm

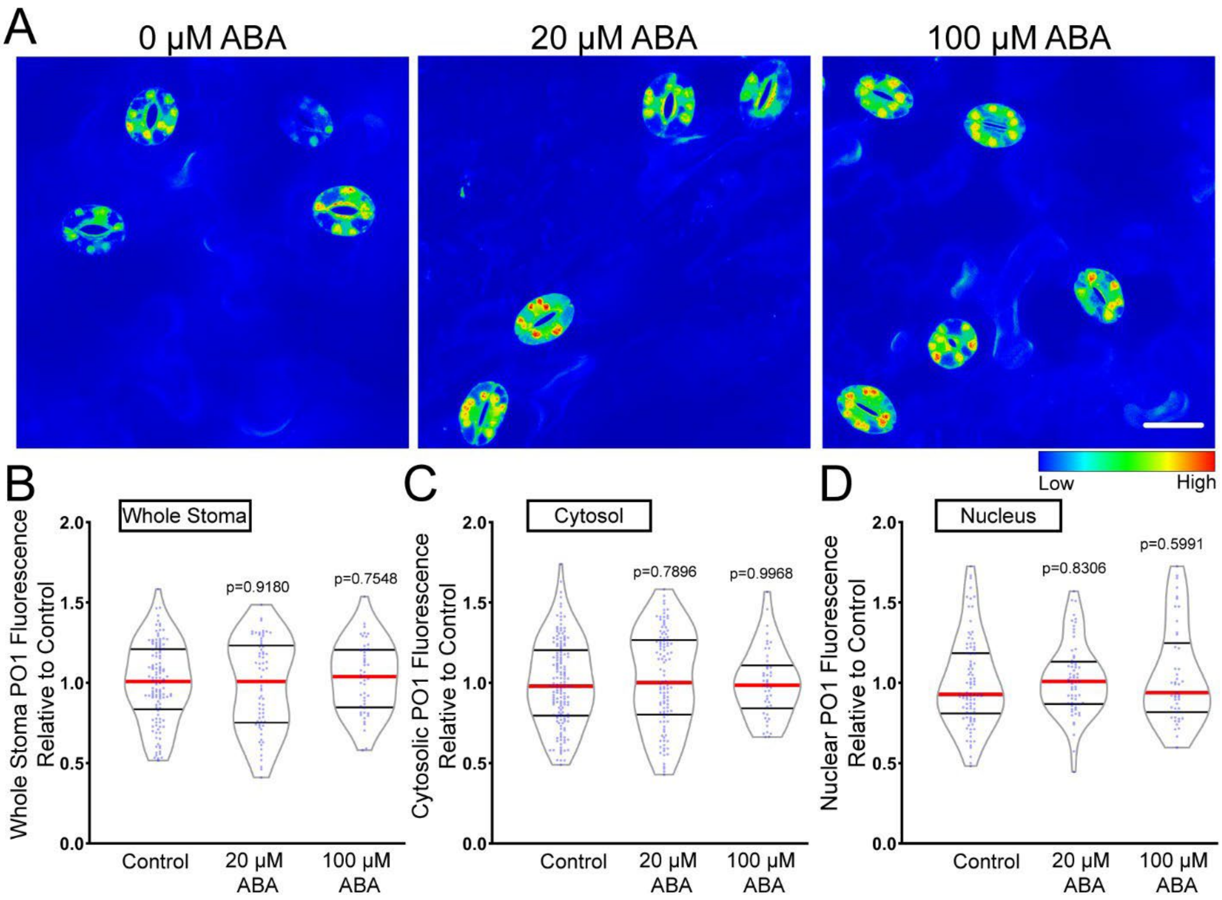


**Supplemental Figure 3.** There are no significant ABA-induced increases in fluorescence of Peroxy Orange 1 in whole stoma, the cytosol, or the nucleus.

A) Confocal micrographs of PO1 signal converted to Lookup Tables (LUT) after treatment with 20 µM or 100 µM ABA or buffer control. Maximum intensity projection of full z-stack is shown. Scale bar: 20µm. Quantifications of PO1 fluorescence in the B) whole stoma, C) cytosol, or D) nucleus. All individual values were normalized to the average of the control treatment for each subcellular location and are displayed on the graph as blue dots with the median shown in red and lower and upper quartiles indicated in black. All p-values were calculated by one-way ANOVA followed by a Tukey's multiple comparisons test from at least three separate experiments (chloroplast n>140, nuclear n>50, cytosol n>50, and puncta n>148).

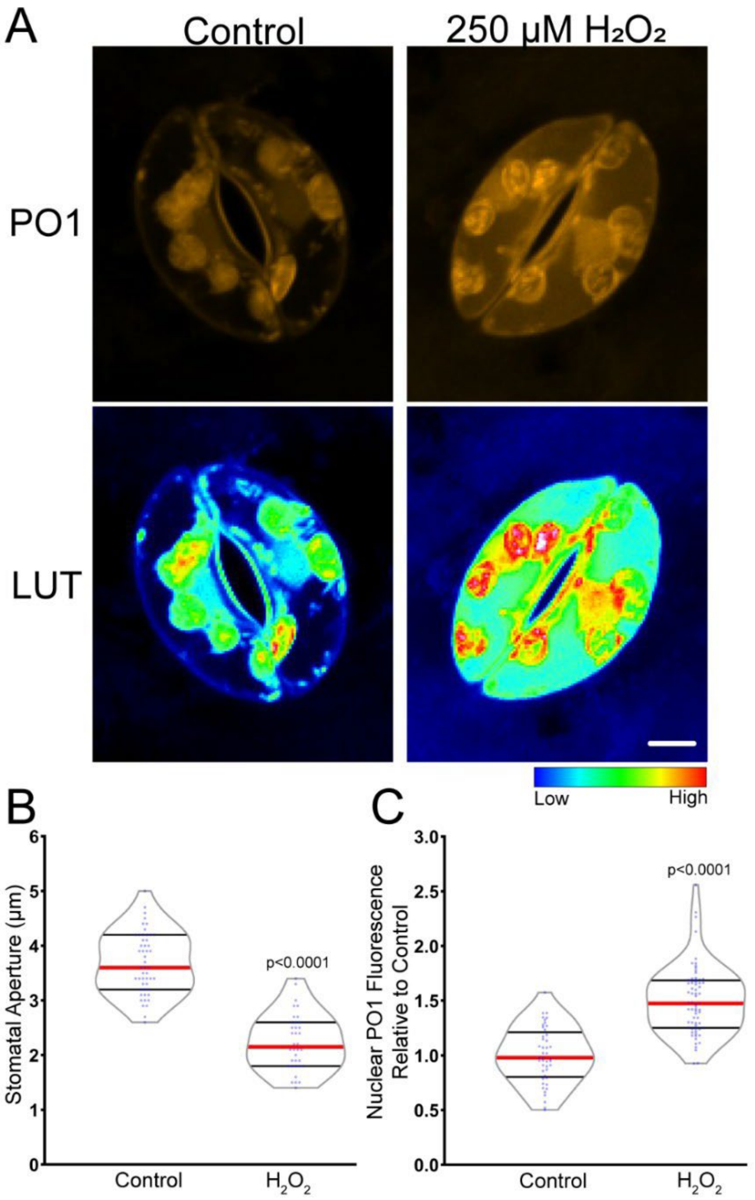


**Supplemental Figure 4.** Treatment with hydrogen peroxide leads to decreased stomatal aperture and elevated PO1 fluorescence.

A) Confocal micrographs of PO1 fluorescence or PO1 fluorescence converted to Lookup Tables (LUT) of Col-0 guard cells treated with buffer control or 250 µM $H_2O_2$. Maximum intensity projection of full z-stack is shown. B) Quantification of stomatal aperture with after treatment with either buffer control of 250 µM $H_20_2$ C) Quantifications of nuclear PO1 fluorescence with $H_2O_2$ treatment or buffer control. All values were normalized to the average of the control treatment and are displayed on the graph as blue dots with the median shown in red and lower and upper quartiles indicated in black. P-values are recorded according to one-way ANOVA followed by Tukey's post hoc test from at least three separate experiments (n>45). Scale bar: 5µm

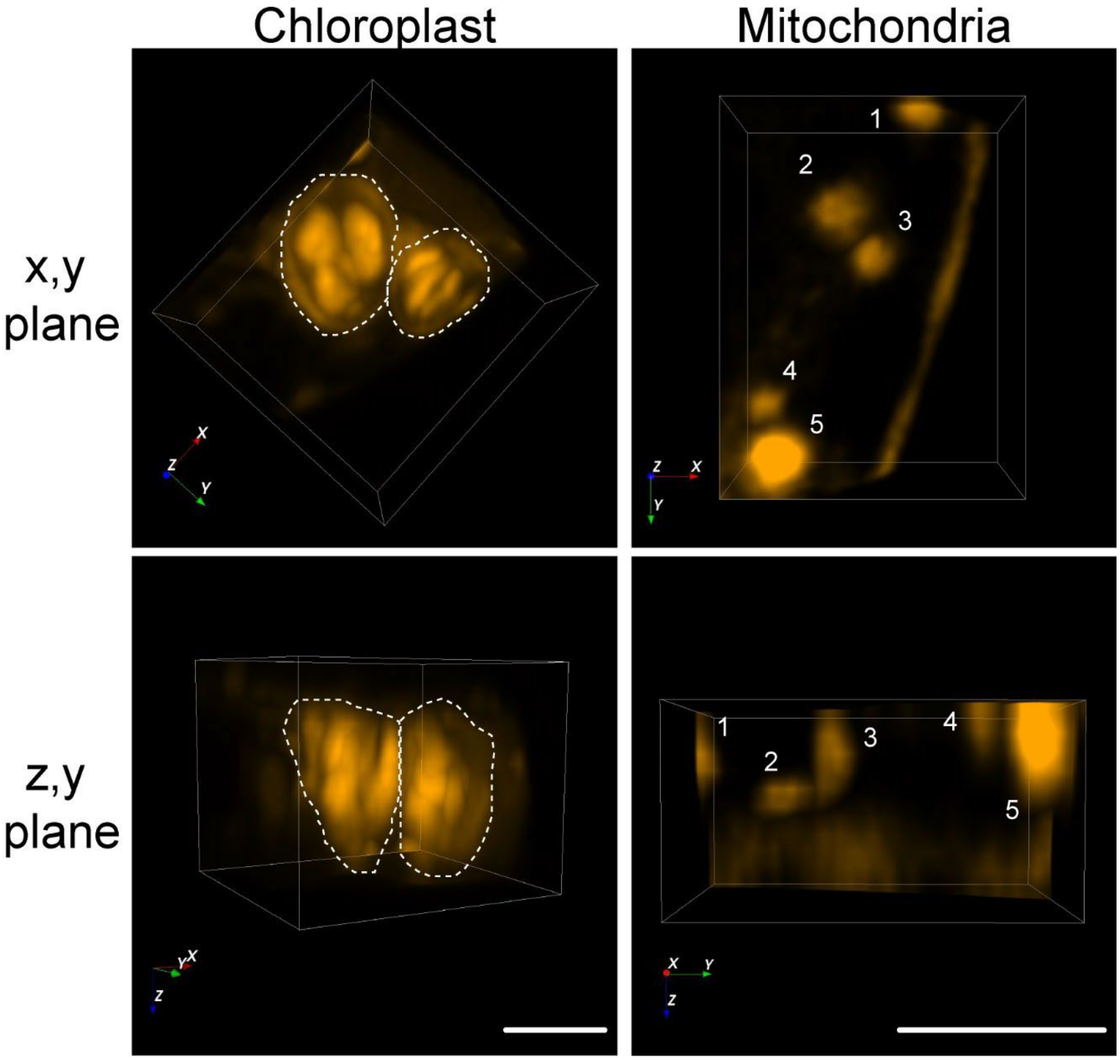


**Supplemental Figure 5.** Mitochondria have uniform PO1 fluorescence, while chloroplasts have localized regions with elevated signal.

3- dimensional renderings of PO1 fluorescence from a guard cell section displaying either a chloroplast or mitochondria. Renderings are oriented in (x,y) plane or (z,y) plane to show depth. Scale bars: 3µm.

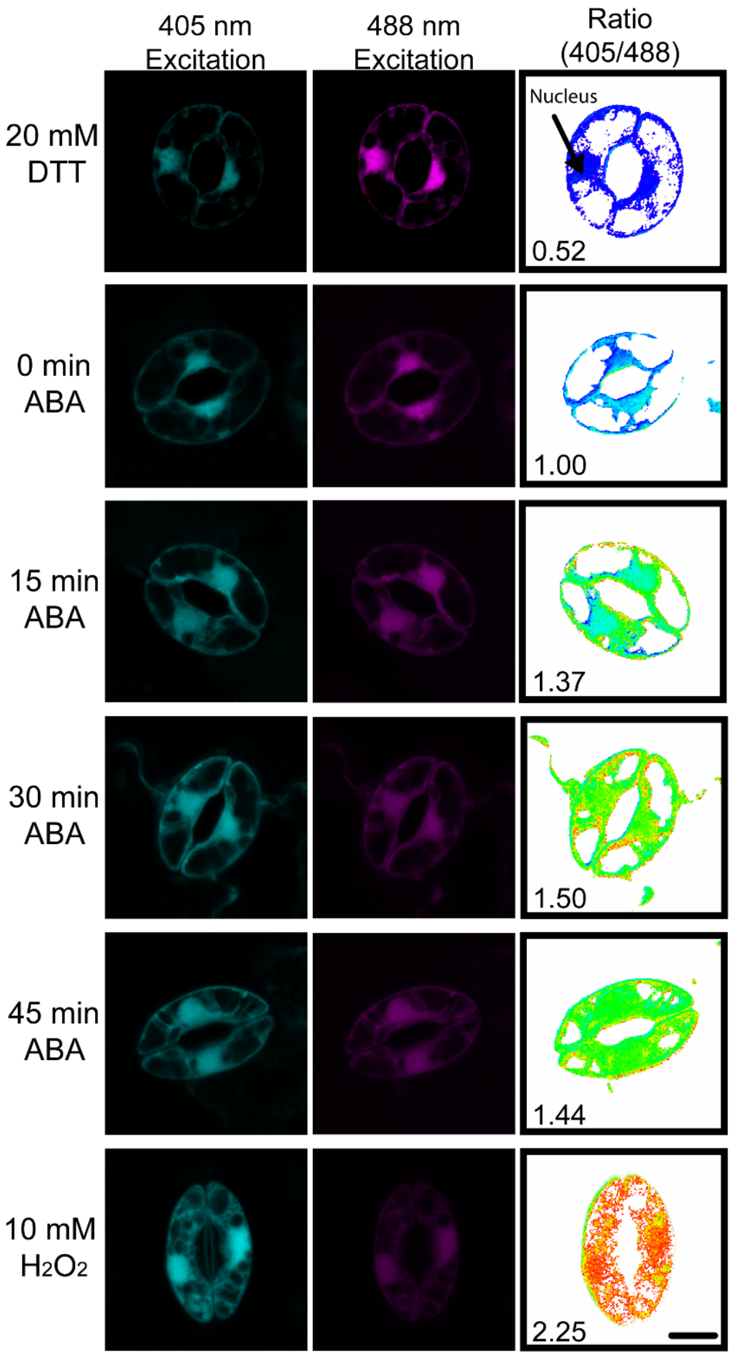


**Supplemental Figure 6.** ABA treatment alters fluorescence of roGFP2-Orp1 in guard cells consistent with elevated hydrogen peroxide. increased.

Representative micrographs of individual channels used to generate ratiometric images (405 nm/488 nm ratio) seen in Figure 4. Under reducing conditions, signal intensity is high following excitation with the 488 nm laser line (magenta) while excitation with the 405 nm laser lines yields low signal intensity (cyan). As roGFP2- Orp1 becomes oxidized, excitation with the 405 nm laser results in increased signal intensity while excitation with the 488 nm line yields decreased fluorescence. Ratiometric images display fluorescence ratios calculated from images taken using sequential excitation at 488 nm and 405 nm for each treatment group. Scale bars: 5µm.

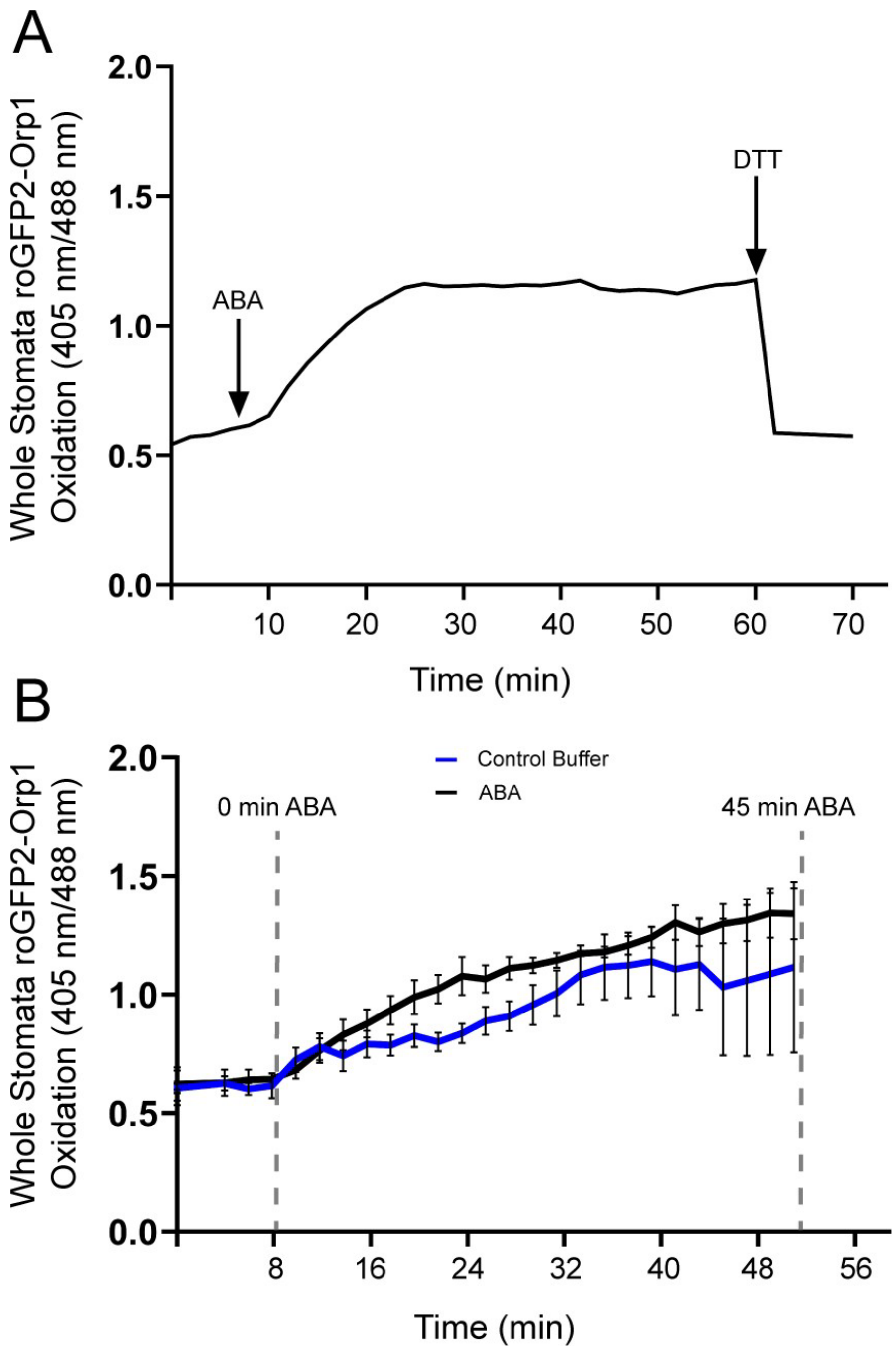


**Supplemental Figure 7.** Time course imaging of the same subset of roGFP2-Orp1 stomata can result in sensor oxidation even under control conditions.

A) Time course showing ABA-driven roGFP2-Orp1 oxidation following treatment with ABA for 50 minutes followed by re-reduction by DTT. B) Time course showing temporal dynamics of roGFP2-Orp1 oxidation following treatment with ABA (black line) or control buffer (blue line) for 45 min. Stomata were imaged under control conditions for a short period before control buffer was replaced with either ABA or fresh control buffer (first dashed line). Images were taken every two minutes for the next 45 minutes (second dashed line).

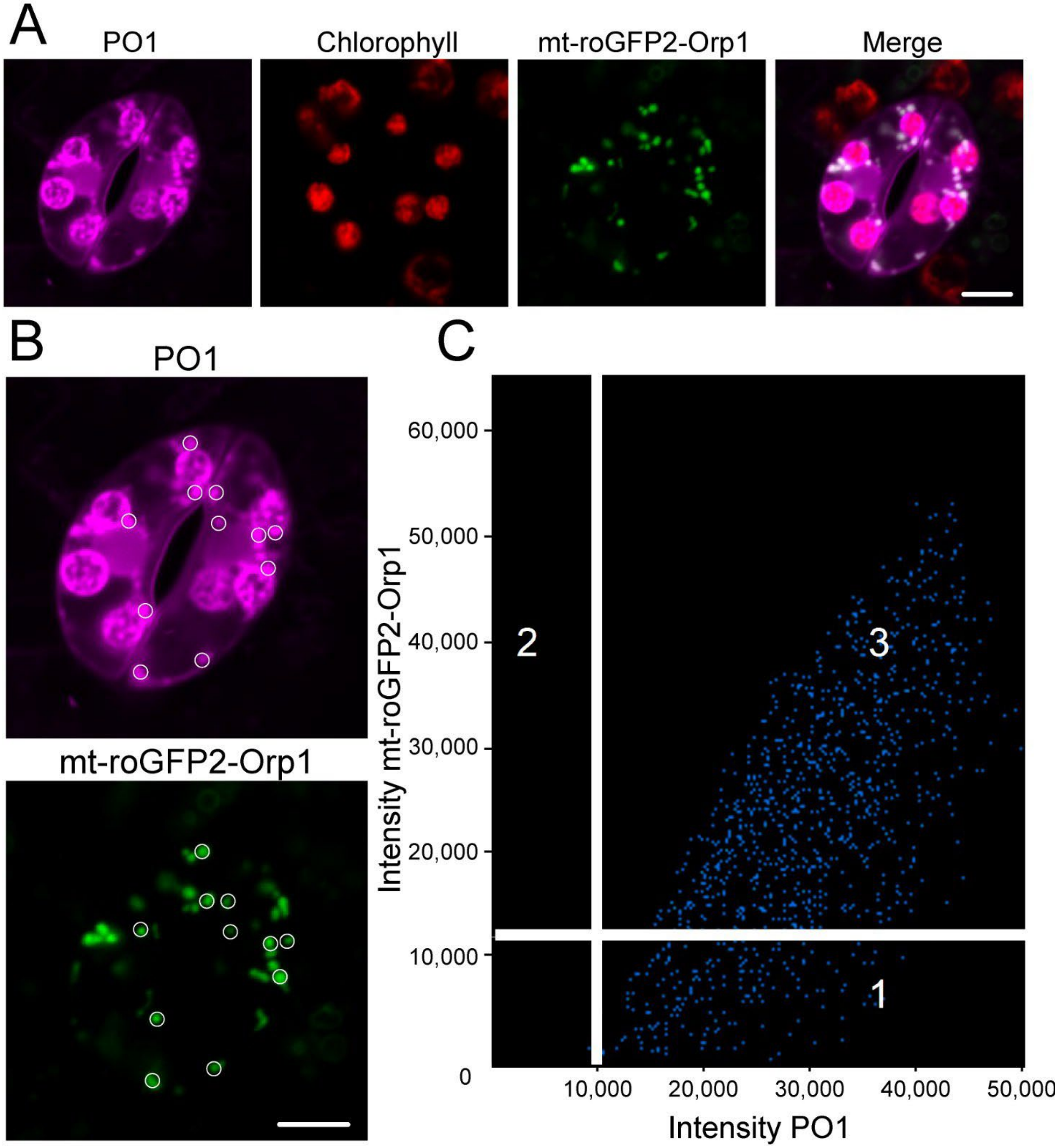


**Supplemental Figure 8.** PO1 and mt-roGFP2-Orp1 colocalize consistent with ABA induced ROS in mitochondria.

A) Confocal micrographs of PO1 fluorescence (magenta), chlorophyll autofluorescence (red), and mt-roGFP2-Orp1 (green), and a merged image showing PO1 colocalized with mt-roGFP2-Orp1 (white). An individual optical slice is shown. B) Regions of interest used to generate weighted colocalization coefficient are circled in white, highlighting PO1 fluorescence colocalizing with mt-roGFP2-Orp1 fluorescence. C) Colocalization graph generated with the ZEN Black colocalization module from regions of interest highlighting mitochondria. Scale bars: 5μm.

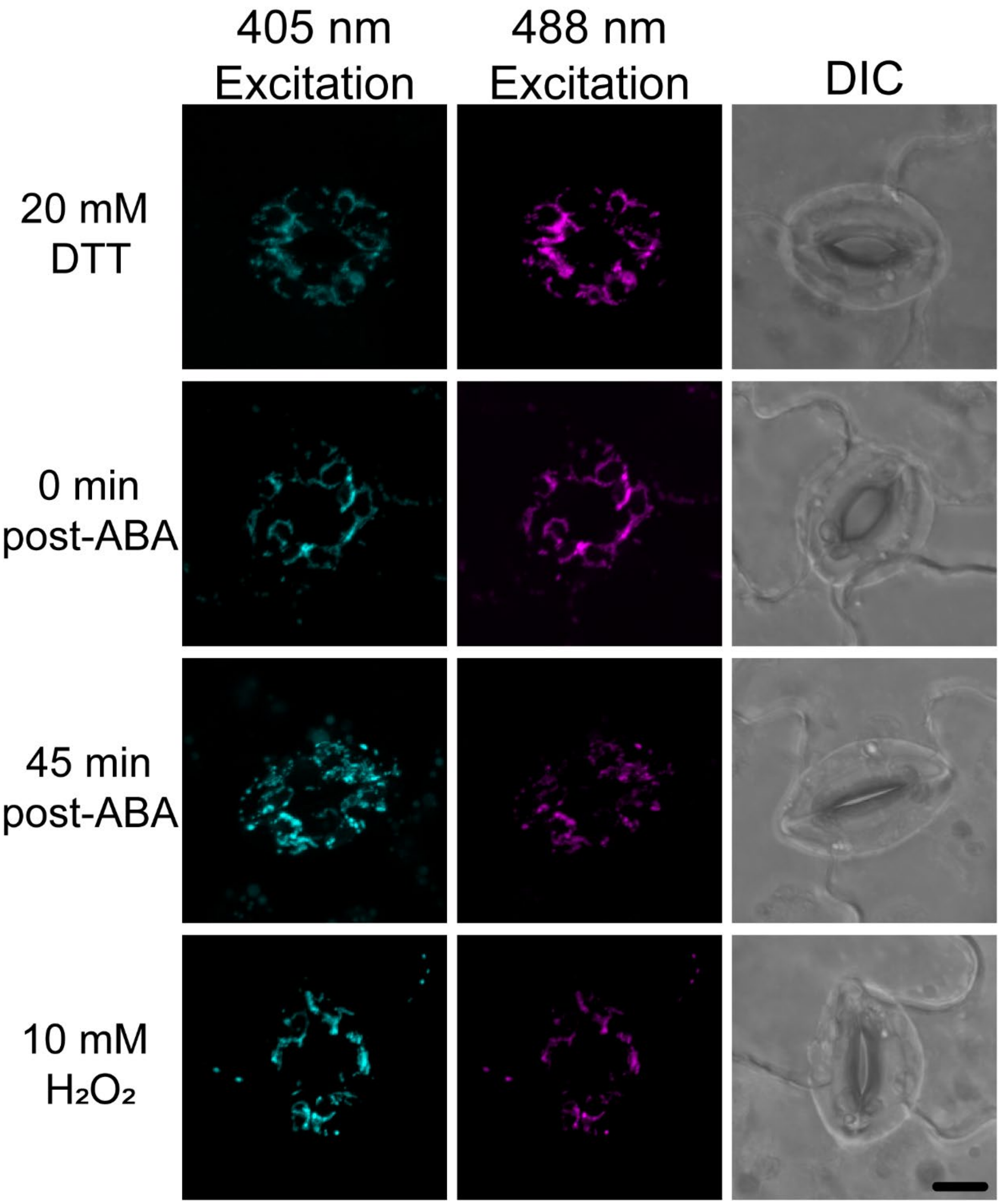


**Supplemental Figure 9.** A 45-minute ABA treatment results in an increased oxidation ratio of mt-roGFP2-Orp1.

Representative micrographs of individual channels used to generate ratiometric images (405 nm/488 nm ratio) seen in Figure 5. Under reducing conditions, signal intensity is high following excitation with the 488 nm laser line (magenta) while excitation with the 405 nm laser lines yields low signal intensity (cyan). As mt-roGFP2-Orp1 becomes oxidized, excitation with the 405 nm laser results in increased signal intensity while signal intensity decreases following excitation with the 488 nm line. Scale bars: 5μm.

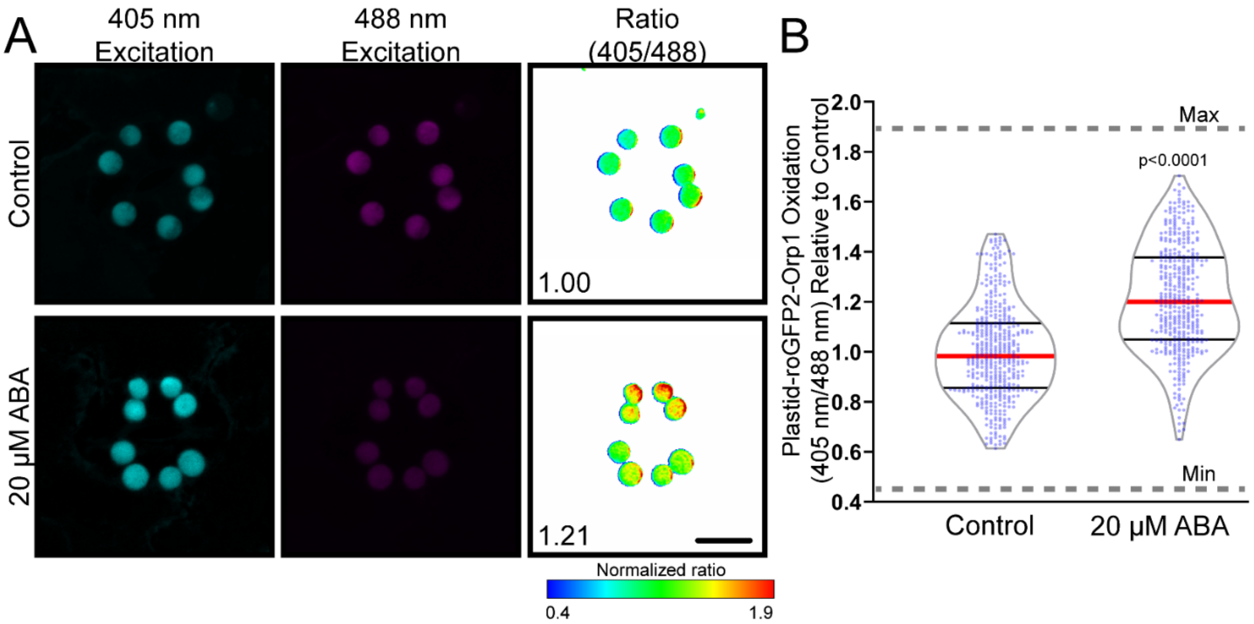


**Supplemental Figure 10.** ABA treatment increased oxidation of plastid-roGFP2-Orp1.

A) Representative micrographs of individual channels as well as ratiometric images of oxidation ratio generated through dividing emission intensity follow excitation at 405 nm by emission intensity after excitation at 488 nm. B) Quantification of plastid-roGFP2-Orp1 ratio in the chloroplasts of guard cells following 45 min ABA treatment. All individual values were normalized to the average of the buffer control treated samples and are displayed on the graph as blue dots with the median shown in red and lower and upper quartiles indicated in black. Minimum and maximum sensor oxidation are shown by treatment with 20 mM DTT or 10 mM $H_2O_2$, respectively. Data are from three separate experiments (n>400 chloroplasts/value). Minimum and maximum sensor oxidation is represented on graphs by gray dashed lines. Listed p-values were determined by one-way ANOVA followed by Tukey's post hoc test. Scale bars: 5µm.

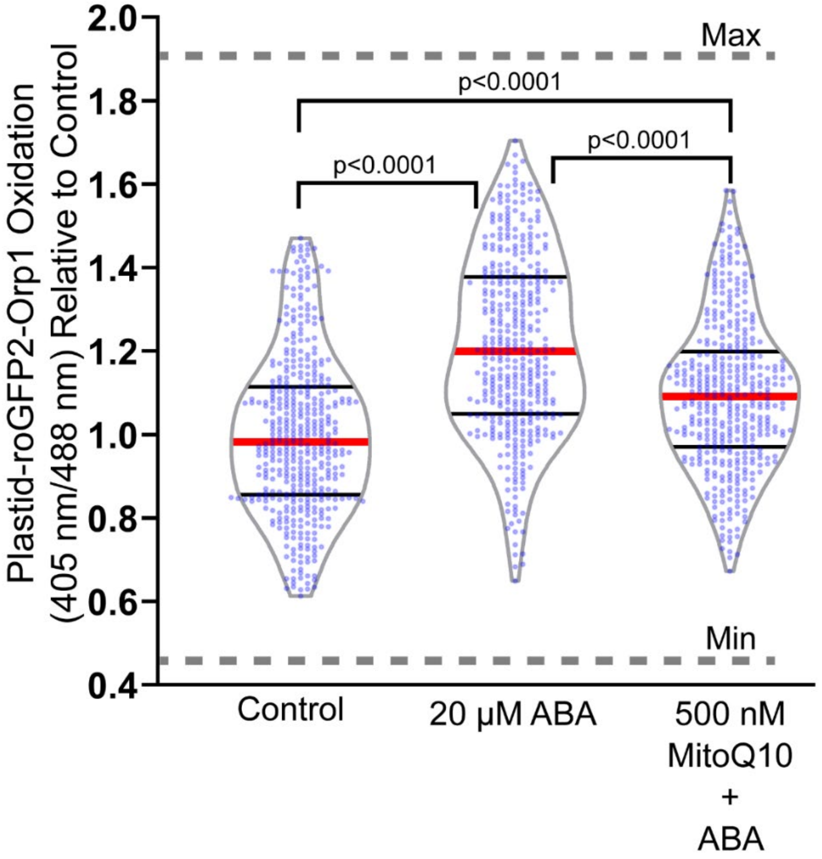


**Supplemental Figure 11.** The ABA-induced oxidation of plastid-roGFP2-Orp1 is reduced by pretreatment with MitoQ.

Quantification of plastid-roGFP2-Orp1 ratio in the chloroplasts of guard cells pretreated with control buffer or 500 nM MitoQ for 3 hours, followed by treatment with 20 μM ABA for 45 min. All individual values were normalized to the average of control treated chloroplasts and are displayed on the graph as blue dots with the median shown in red and lower and upper quartiles indicated in black. Minimum and maximum sensor oxidation are shown by treatment with 20 mM DTT or 10 mM $H_2O_2$, respectively. Data are reported from three separate experiments (n>400 chloroplasts). Minimum and maximum sensor oxidation is represented on graphs by gray dashed lines. Listed p-values were determined by one-way ANOVA followed by Tukey's post hoc test. Scale bars: 5μm.

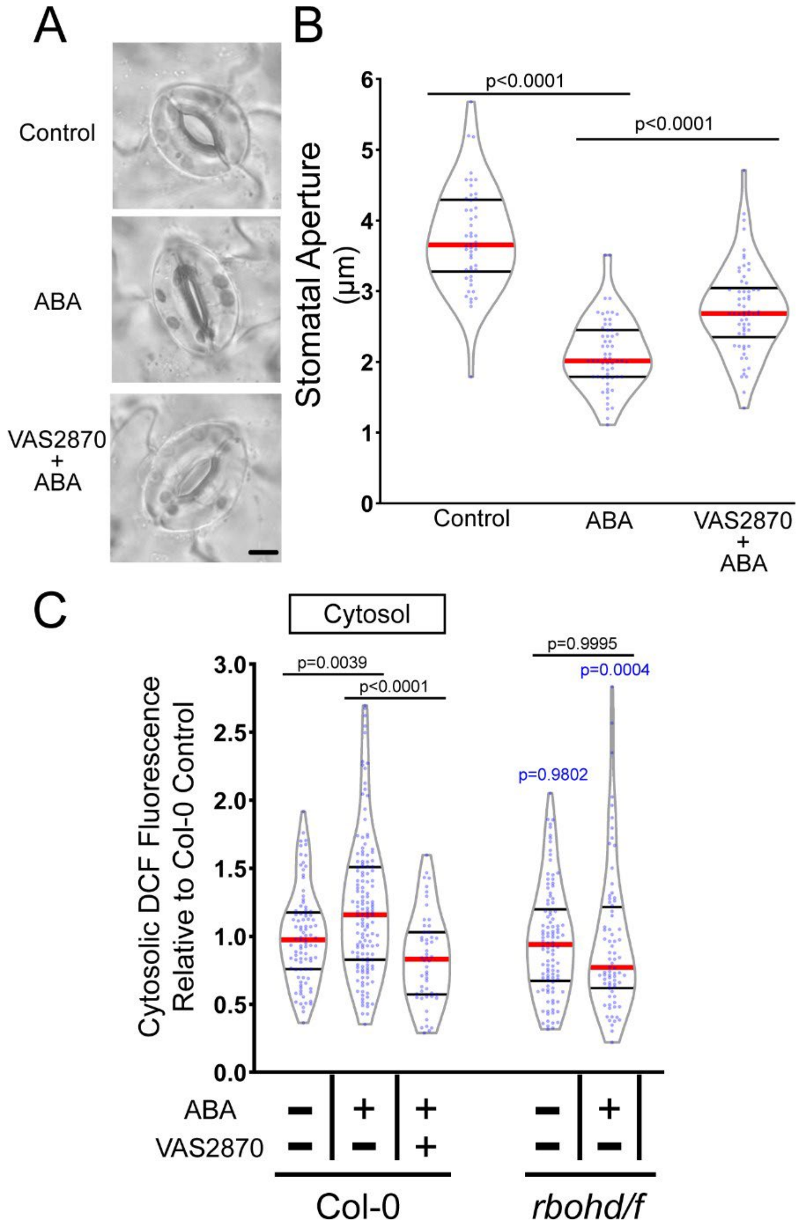


**Supplemental Figure 12.** The RBOH-selective inhibitor VAS2870 reduces ABA-dependent guard cell closure and DCF accumulation.

A) Stomatal apertures of leaves of Col-0 treated with a buffer control, ABA, or pre-incubated with 10 µM VAS2870 for 1 hr followed by ABA treatment for 45 min. B) Stomatal apertures of buffer control, ABA, or VAS2870 + ABA were quantified immediately following treatment. C) Violin plots show quantifications of DCF fluorescence in the cytosol following treatment with control buffer, ABA, or pre-treated with VAS2870 and then treated with ABA from three separate experiments (n>52). All individual values were normalized to the average of the control treatment in Col- 0 for each subcellular location and are displayed on the graph as blue dots with the median shown in red and lower and upper quartiles indicated in black. P-values in black font represent the significance of differences between treatments in the same genotype spanning the compared treatments. P-values in blue font representing the significance of differences between the indicated genotypes and Col-0. P-values are recorded according to two-way ANOVA followed by Tukey's post hoc test.
